# Role of YAP in early ectodermal specification and a Huntington’s Disease model of human neurulation

**DOI:** 10.1101/2021.08.11.455964

**Authors:** Francesco M. Piccolo, Nathaniel R. Kastan, Tomomi Haremaki, Qingyun Tian, Tiago L. Laundos, Riccardo De Santis, Thomas S. Carroll, Ji-Dung Luo, Ksenia Gnedeva, Fred Etoc, A. J. Hudspeth, Ali H. Brivanlou

## Abstract

The Hippo pathway, a highly conserved signaling cascade that functions as an integrator of molecular signals and biophysical states, ultimately impinges upon the transcription coactivator Yes-associated protein 1 (YAP). Hippo-YAP signaling has been shown to play key roles both at the early embryonic stages of implantation and gastrulation, and later during neurogenesis. To explore YAP’s potential role in neurulation, we used self-organizing neuruloids grown from human embryonic stem cells on micropatterned substrates. We identified YAP activation as a key lineage determinant, first between neuronal ectoderm and non-neuronal ectoderm, and later between epidermis and neural crest, indicating that YAP activity can enhance the effect of BMP4 stimulation and therefore affect ectodermal specification at this developmental stage. Because aberrant Hippo-YAP signaling has been implicated in the pathology of Huntington’s Disease (HD), we used isogenic mutant neuruloids to explore the relationship between signaling and the disease. We found that HD neuruloids demonstrate ectopic activation of gene targets of YAP and that pharmacological reduction of YAP’s transcriptional activity can partially rescue the HD phenotype.

## Introduction

Although the study of embryogenesis benefits from a broad array of model organisms and the conservation of mechanisms across species, elucidation of the particularities of human development remains challenging. The use of human pluripotent stem cells for the creation of synthetic human organoids has greatly enhanced our ability to investigate human development. For example, the process of neurulation can be mimicked *in vitro* through micropattern-based neuruloids (Haremaki et al., 2019). This technique, which allows the generation of hundreds of virtually identical organotypic cultures, offers a powerful opportunity to investigate the molecular and biophysical principles of human neurulation and the associated diseases.

The Hippo pathway is an ancient and highly conserved signaling cascade that operates in numerous cell types and a variety of organisms. In contrast to developmental pathways that require specific ligand-receptor interactions, the Hippo pathway functions as an integrator of molecular signals and biophysical states, including cell polarity, adhesion, GPCR signaling, and mechanical forces sensed through the cytoskeleton. With key roles in development, homeostasis, and regeneration, the pathway interacts with other developmental and regulatory signals including Wnt, Notch, and TGFβ. Despite the apparent complexity, the core cascade is relatively simple, including the parallel mammalian STE20-like kinases 1 and 2 (MST1/2) and mitogen-activated protein kinase kinase kinase kinase (MAP4K). The activity of these enzymes leads to phosphorylation of large tumor suppressor kinases 1 and 2 (LATS1/2), which in turn phosphorylate and reduce the nuclear flux of Yes associated protein 1 (YAP) and the homologous WW domain-containing transcription regulator 1 (TAZ). When they translocate to the nucleus, YAP and TAZ cooperate with TEA-domain transcription factors (TEAD1/2/3/4) to activate the expression of a variety of genes associated with proliferation, cell survival, de-differentiation, and cellular morphogenesis (Moya & Halder, 2019).

Hippo-YAP signaling is important at various embryonic stages, including in zygotes and blastomeres, the inner cell mass (ICM), trophectoderm (TE), and later the heart, brain, eye, lung, and kidney. Because Yap^−/−^ mice die at E8.5 owing to failure of endothelium formation in the yolk sac, and Yap^−/−^,Taz^−/−^ embryos fail prior to the morula stage, investigating the potential role of Yap in neurulation *in vivo* requires complex transgenic strategies (Wu & Guan, 2021). The use of neuruloids derived from human embryonic stem cells (hESCs) therefore offers a convenient, highly reproducible, and quantitative alternative means of investigating the role of Hippo signaling in neurulation.

Here we identify YAP activation as a key lineage determinant, first between neuronal ectoderm (NE) and non-neuronal ectoderm (NNE), and later between epidermis and neural crest. Exploiting the reproducible phenotype of Huntington’s Disease (HD) mutations in neuruloids, we additionally find that HD neuruloids demonstrate stronger YAP’s downstream transcriptional signature and that the pharmacological reduction of YAP’s transcriptional activity partially reverses this effect.

## Results

### YAP activity differs between neural and non-neural ectoderm

In order to study the role of the Hippo pathway during human neurulation *in vitro*, we applied a neuruloid assay (Haremaki et al., 2019). After seeding on circular, 500 μm-diameter islands, hESCs underwent neural induction as a result of dual SMAD inhibition for three days (SB431542 and LDN193189, SB+LDN) followed by BMP4 treatment (SB+BMP4). Because BMP4 receptivity was limited to the edges of the colonies (Haremaki et al., 2019), non-neuronal ectoderm (NNE) emerged there by the fourth day—which we designate hereafter as D4—and neuronal ectoderm (NE) remained unperturbed in the center (Figure 1A).

**Figure 1.**
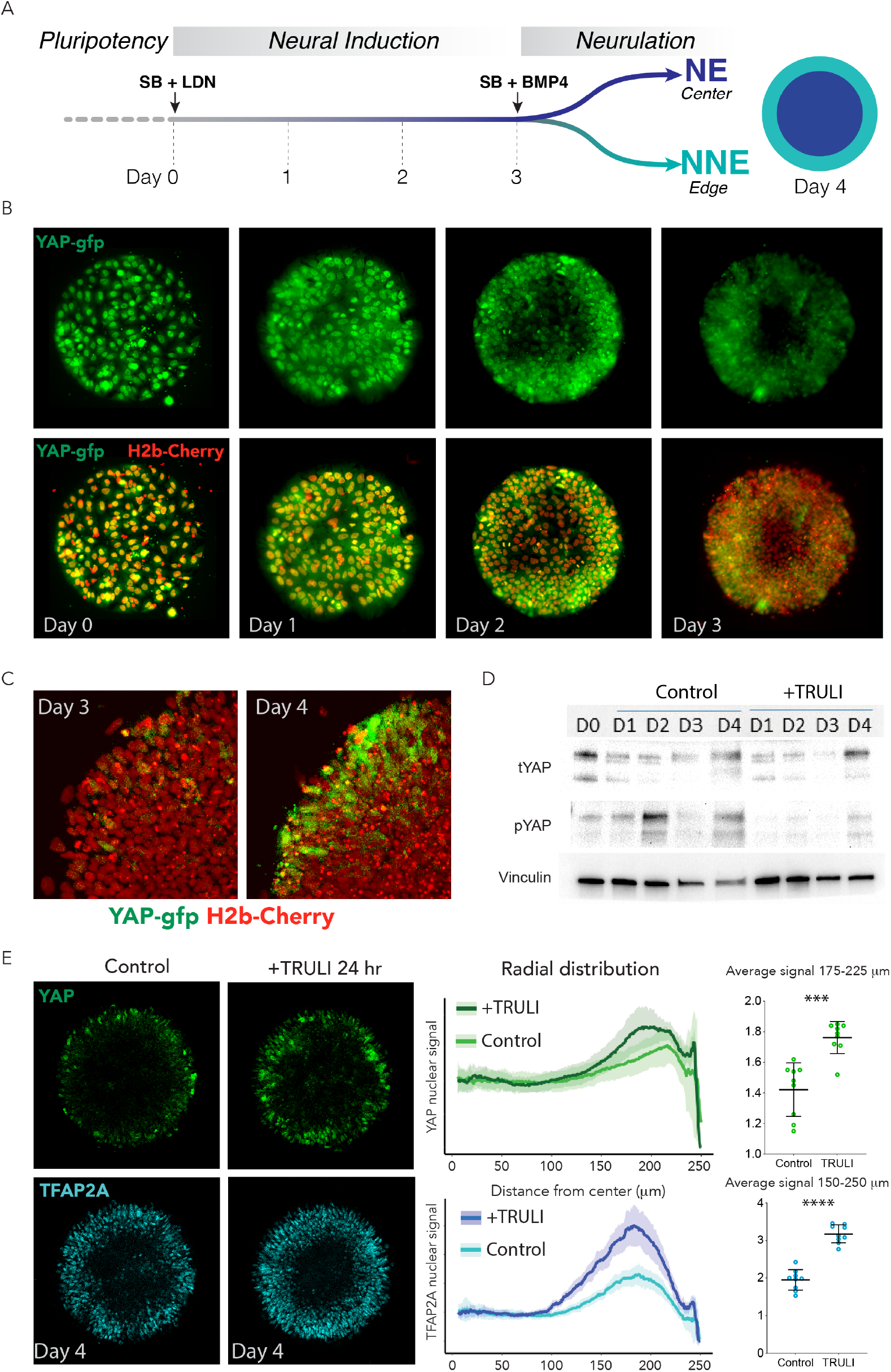
Dynamics of YAP expression and localization in early neuruloids. A) During the first four days of the differentiation protocol, pluripotent hESCs seeded on a micropattern undergo three days of neural induction (SB+LDN) followed by BMP4 induction of neurulation (SB+BMP4). B) Fluorescent images of living YAP-GFP and H2b-Cherry hESC colonies during neural induction show the progressive shift of YAP from nuclei into cytoplasm and a reduction in expression level. C) Fluorescent images portray one quadrant of a colony acquired before (D3) and 24 hrs after (D4) BMP4 administration. D) Immunoblots from D0 to D4 of the neuruloid protocol illustrate the decrease in YAP phosphorylated at residue S127 (pYAP) upon treatment with 10 μM TRULI. The amount of total YAP protein (tYAP) initially declines but recovers by D4. Vinculin serves as a loading control. E) Left: Immunolabeling of D4 neuruloids demonstrates the increased concentration of YAP and TFAP2A, a marker of NNE, at the edges of colonies treated for 24 hours with 10 μM TRULI. Right: Radial distribution of YAP and TFAP2A nuclear signals from several micropattern colonies quantitate the result. The average values at the edge of each colony are also shown. ****, *p* < 0.0001; ***, *p* < 0.001 in unpaired *t*-tests comparing untreated (control) and TRULI-treated samples.

To assess Hippo-YAP signaling during these early stages of neuruloid formation, we genetically engineered the RUES2 hESC line (National Institutes of Health #0013) to endogenously and biallelically tag the C-terminus of YAP protein with green-fluorescent protein (GFP; Supplemental Figures S1 and S2; Franklin et al., 2020). During the pluripotency state at the outset of culture, D0, YAP was highly expressed and occurred within nuclei, but upon neural induction YAP was progressively enriched in the cytoplasm and by D3 was downregulated to a low level of expression (Figure 1B,C).

Twenty-four hours after BMP4 stimulation, live-imaging experiments on D4 revealed that YAP expression was upregulated at the edge of the micropattern colony, in the region where the NNE lineage would emerge. Near the border of the colony, YAP could be observed to undergo sporadic nuclear-cytoplasmic flux (Figure 1C; Supplemental Movie 1) that suggested active regulation of the protein’s location in the developing NNE lineage.

### YAP activation with TRULI augments differentiation of non-neural ectoderm

Immunoblot analyses of D4 neuruloids confirmed a rise in YAP levels and revealed phosphorylation of YAP at residue S127, consistent with regulation through the Hippo pathway (Figure 1D). Concomitant application of the Lats-kinase inhibitor TRULI (Kastan et al., 2021) suppressed YAP phosphorylation without altering the dynamics of the protein’s concentration during this initial phase of neuruloid self-organization (Figure 1D). Consistent with the loss of S127 phosphorylation, administration of TRULI in conjunction with BMP4 increased the level of YAP protein and induced its nuclear localization in cells at the edges of neuruloid colonies (Figure 1E; Supplemental Figure 2A, B; Supplemental Movie 2). Finally, TRULI-treated neuruloids demonstrated more robust TFAP2A induction, suggestive of augmented NNE specification (Figure 1E). These results together suggest that YAP expression is upregulated in response to BMP4 at the periphery of the colony, is subsequently regulated in part by the Hippo pathway, and contributes directly to induction of the NNE lineage.

**Figure 2.**
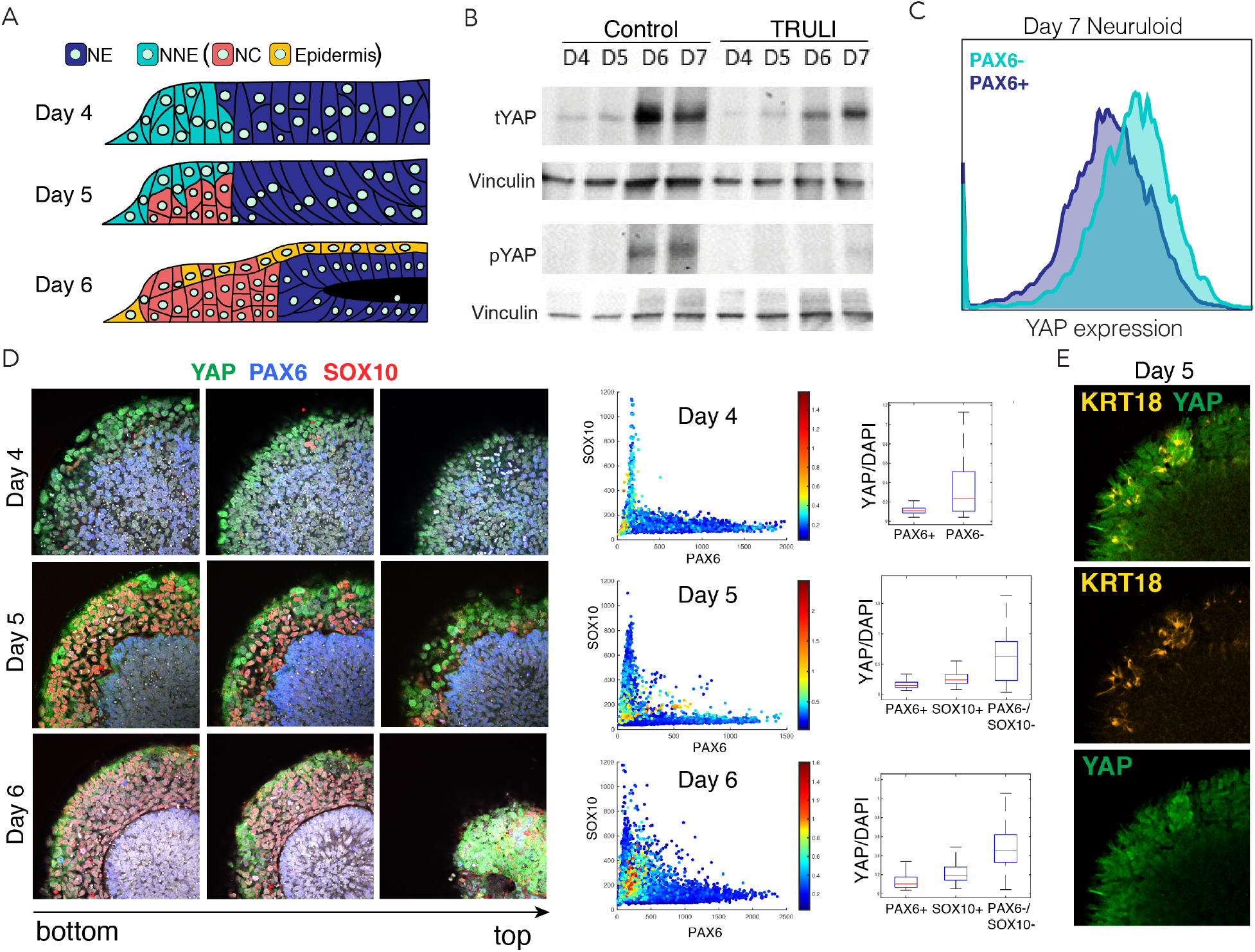
Dynamics of YAP expression and localization in late neuruloids. A) In a schematic side view of the final phase of neuruloid formation, neural ectoderm (NE, dark blue) at the center is surrounded by non-neural ectoderm (NNE, cyan), which subsequently differentiates into neural crest (NC, red) and epidermis (yellow). (B) Immunoblot analysis of neuruloids at D4-D7 demonstrates the suppression of YAP phosphorylated on residue S127 (pYAP) by 10 μM TRULI. The concentration of total YAP protein (tYAP) increases under both conditions. Vinculin provides a loading control. C) Analysis by fluorescence-activated cell sorting of D7 neuruloids shows less expression of total YAP protein in NE (PAX6+) than in NNE (PAX6−). D) Immunohistochemistry during the late phase of neuruloid formation, D4-D6, shows the progressive decline in nuclear YAP in NE and NC lineages. For four neuruloids at each time, the nuclear signals of YAP, PAX6, and SOX10 were determined in individual cells. Scatterplots and plots of normalized YAP signals quantify the effect. E) Immunofluorescence images of one quadrant of a D5 neuruloid demonstrate the presence of KRT18, a label for epidermis, in cells enriched for nuclear YAP.

### YAP activity contributes to differentiation of neural crest and epidermis

After the initial separation of the NE and NNE lineages at D4, maturation to a complete neuruloid occurs by D7 (Haremaki et al., 2019). In the course of this process, the NE organizes into a compact, closed structure at the center of the colony. The NNE differentiates further, forming two principal derivatives: neural crest (NC), which is found in a radially symmetric arrangement at the periphery of the neuruloid, and epidermal cells (E), which grow atop the culture (Figure 2A). The overall structure models the early stages of human neurulation (Haremaki et al., 2019). On D6, YAP levels rose sharply with a concomitant increase of phospho-S127 YAP (Figure 2B). Fluorescence-activated cell sorting on D7 demonstrated that enrichment of YAP in NNE (PAX6−) relative to NE (PAX6+) persisted through maturation of the neuruloid (Figure 2C).

We assessed the expression and localization of YAP within the major ectodermal lineages at different times during neuruloid formation (Figure 2D, E). We confirmed the exclusion of YAP from the nuclei both of NE (PAX6+) cells and of the emerging neural-crest cells (SOX10+; Figure 2D). YAP was most notably present in the nuclei of cells expressing the early epidermal marker KRT18 (Figure 2E).

Given that YAP is enriched first in NNE and subsequently in epidermis, we inquired whether YAP signaling stimulates their differentiation. Taking advantage of a protocol for the direct differentiation of neural crest (Tchieu et al., 2017), we produced SOX10+ colonies with only sparse KRT18+ cells (Figure S3A). The administration of TRULI, which enhanced YAP activity (Figures 1E and S2), resulted in the complete loss of NC− like SOX10+ colonies and a significant increase in KRT18 expression (Figure S3A). Consistent with these observations, in a converse experiment TRULI treatment boosted KRT18 expression in an epidermal-differentiation protocol (Figure S3B; Tchieu et al., 2017). Furthermore, TRULI-treated neuruloids displayed increased differentiation toward epidermis and a reduction in SOX10+ cells (Figure 3). Surprisingly, TRULI treatment also precluded the closure of the central PAX6+ structure, the simulacrum of the neural tube, despite its being devoid of detectable YAP (Figure 3).

**Figure 3.**
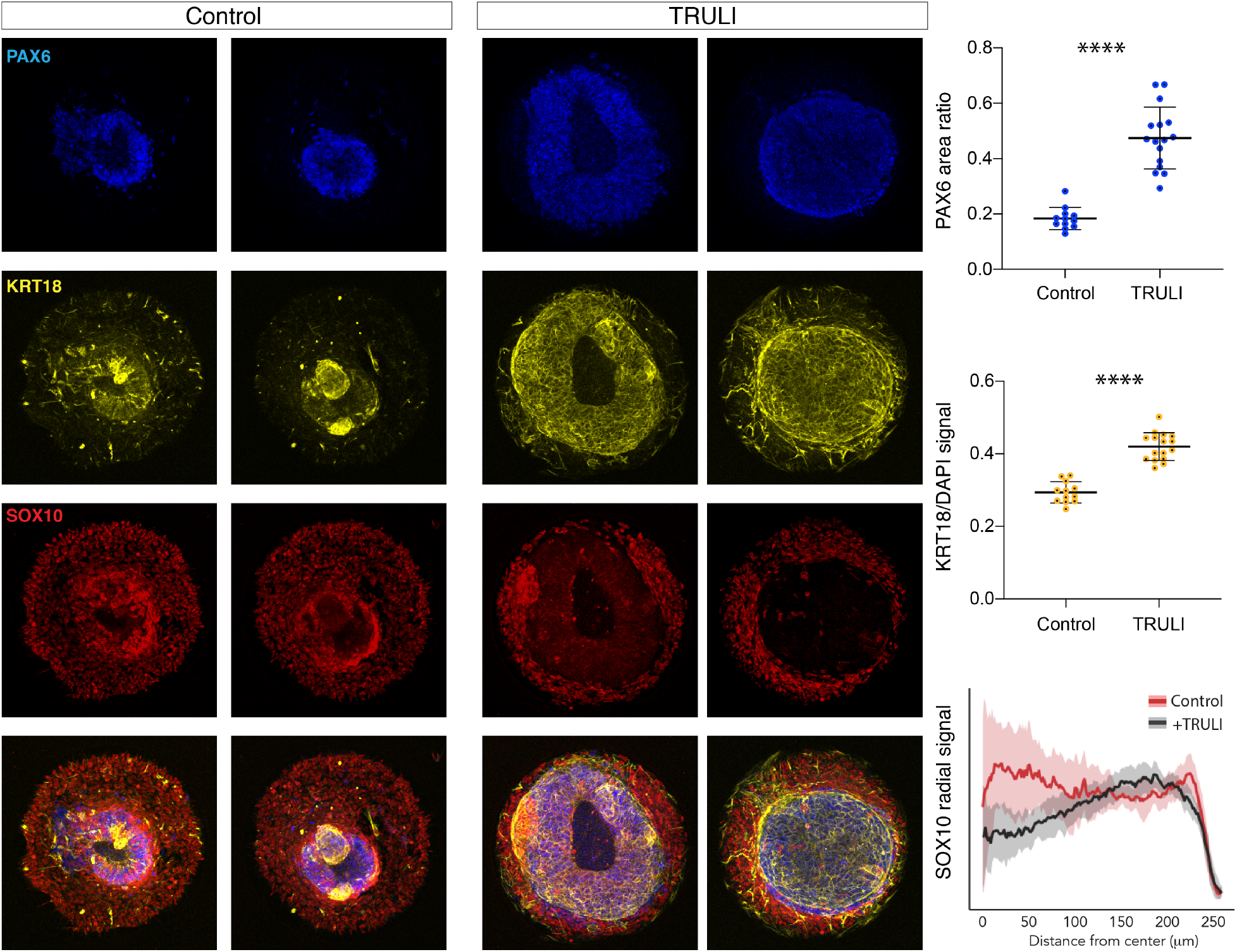
Effect of sustained YAP activation on neural crest and neural epithelium. Immunofluorescence images of D7 WT neuruloids demonstrate that treatment with 10 μM TRULI enhances the expression of neural ectoderm marked by PAX6 and of epidermis marked by KRT18 at the expense of NC labeled by SOX10. The plots quantify the fractions of the areas of several colonies marked by PAX6 and KRT18 (as normalized to DAPI), as well as the radial distribution of SOX10. ****, *p* < 0.0001 in an unpaired *t*-test comparing untreated (control) and TRULI-treated samples.

These experiments highlight the role of YAP in an *in vitro* model of human neurulation and suggest that the protein is dynamically regulated through canonical Hippo signaling during the specification of ectodermal lineages. Following an initial downregulation in ectodermal cells, the expression of YAP protein is induced by BMP4 stimulation and its activity supports the induction of the NNE lineage. YAP subsequently contributes directly to the differentiation of NNE by repressing the development of the NC lineage and promoting the emergence of epidermal cells.

### Developing HD neuruloids display increased YAP activity

The development of HD can be modeled in human neuruloids (Haremaki *et al*., 2019). Using this approach, we characterized the phenotype of neuruloids made from hESCs bearing 56 CAG repeats, an expansion characteristic of HD. Hereafter we term these HD neuruloids, as opposed to wild-type (WT) neuruloids made from the original, isogenic RUES2 cell line. Like those treated with TRULI, HD neuruloids failed to display closure of the central PAX6+ area (Figure 3). Intrigued by the similarities between the two phenotypes, and in view of the potential roles of YAP in human ectodermal differentiation, we next investigated whether the HD phenotype in neuruloids stems from hyperactivity of YAP.

To directly compare the behavior of YAP in WT and HD neuruloids, we genetically engineered an HD line to express endogenously tagged YAP-GFP (Figures S1 and S2). The behavior of YAP in WT and HD neuruloids was indistinguishable during the first three days of neural induction, when YAP was excluded from nuclei and progressively downregulated (Figure S4). Both WT and HD lines likewise upregulated YAP expression at the edges of colonies upon BMP4 stimulation. However, at D4 YAP appeared to be more enriched in nuclei at the most peripheral region of HD neuruloids compared to WT controls (Figure 4A; Supplemental Movie 3). A similar result was obtained by immunofluorescence microscopy, which confirmed elevated YAP nuclear signals in HD colonies (Figure 4B). Consistent with these observations, immunoblotting analysis of D4 HD neuruloids revealed a reduction in the phosphorylated form of YAP in comparison to WT controls (Figure 4C). These observations suggest increased YAP activity in early HD neuruloids.

**Figure 4.**
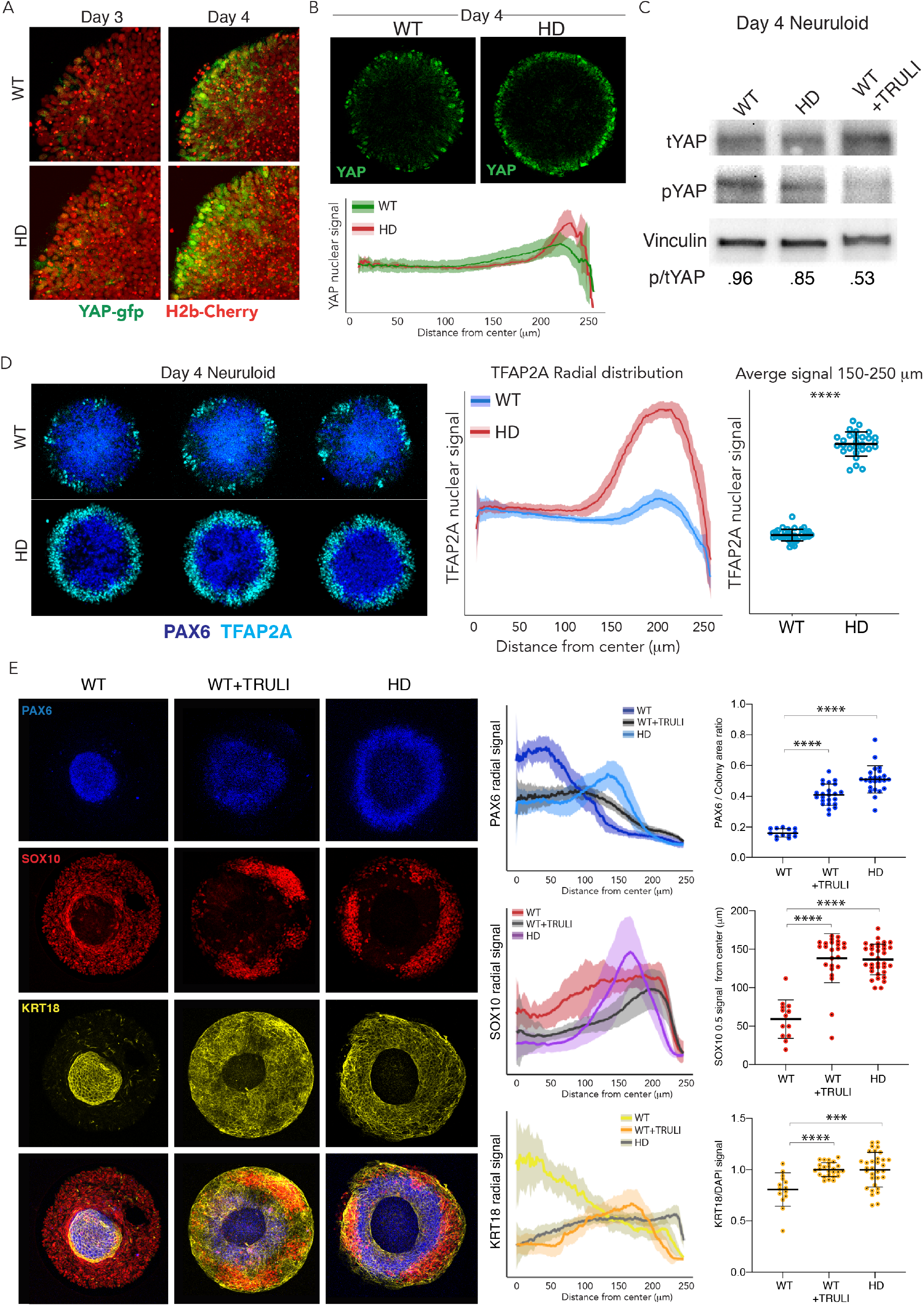
Dysregulation of YAP localization in HD neuruloids. A) Fluorescent images of the same portions of living neuruloids before (D3) and 24 hr after BMP4 administration (D4) show increased nuclear YAP in HD than in WT neuruloids. B) Top, YAP immunofluorescence images of D4 neuruloids confirm the effect. Bottom, the radial distribution of YAP nuclear signals is quantitated for 16 neuruloid colonies for each condition. C) Immunoblot analysis of WT and HD neuruloids at D4 shows reduction of YAP phosphorylated on residue S127 (pYAP) in HD colonies, similar to that after TRULI treatment of WT colonies. Vinculin provides a loading control. D) Left, Immunofluorescence images of D4 neuruloids demonstrate enhanced TFAP2A at the perimeters of HD colonies. Right, quantitative analysis of nuclear TFAP2A signals confirms the effect. ****, *p* < 0.0001 in unpaired *t*-tests comparing WT and HD samples. E) WT neuruloids, WT neuruloids treated with 10 μM TRULI, and HD neuruloids. Left, immunofluorescence images of D7 neuruloids demonstrate the similar phenotypes of TRULI treatment and HD: expansion of NE (PAX6), the diminishment of NC (SOX10), and the enhancement of epidermis (KRT18). Right, quantitative analysis of the three experimental conditions showing radial distributions of the lineage markers PAX6, SOX10, and KRT18 in several colonies. For each colony, plots show the area of the PAX6+ central region as a fraction of the entire colony and the radial distance from the colony’s center to the half-maximal intensity of the SOX10 domain. Plots of the KRT18 signal normalized by DAPI labeling confirm the result. ****, *p* < 0.0001 in unpaired *t*-tests comparing the three experimental conditions.

As observed after TRULI treatment of WT cells (Figure 1E), D4 HD neuruloids displayed more TFAP2A at the edges of colonies (Figures 4D and S5). This observation implies that increased YAP activity in early HD neuruloids has early and direct consequences for fate acquisition in the developing NNE. Consistent with elevated YAP activity, by D7 both HD neuruloids and TRULI-treated WT neuruloids displayed a remarkably expanded epidermal lineage and a reduction in the neural-crest population, as well as a failure in NE compaction (Figure 4E).

These results indicate that neuruloids obtained from hESCs carrying a HD mutation display elevated YAP activity. Consistent with our observation that YAP activity regulates NNE and epidermal formation, at D4 the HD neuruloids demonstrate more robust induction of NNE, which results by D7 in significant expansion of the epidermis at the expense of neural crest.

### Neural ectoderm of HD neuruloids displays ectopic YAP activity

Because we observed elevated YAP activity as early as D4, we characterized this intermediate state by single-cell transcriptomic analysis. The expression profiles of lineage-marker genes in this dataset allowed us to annotate the various cell populations (Figure S6A). As expected, *PAX6* and *TFAP2A* were mutually exclusive and identified NE and NNE, respectively (Figure S6B). The NNE cell cluster could be subdivided into two sub-populations: NC progenitors expressing *FOXD3* and early epidermis expressing *KRT18*. These populations recapitulated the three major human ectodermal lineages: neural ectoderm, neural crest, and epidermis (Figure S6C).

Using the aforementioned markers to identify the progenitors of the major ectodermal lineages, we used scRNA-seq to assess their relative levels of YAP activity. *YAP* and key Hippo pathway components were expressed in all three lineages, most strongly in NC and epidermis (Figures 5A,B and S7). However, the expression levels of YAP target genes, such as CTGF, TAGLN and TUBB6, were much higher in epidermis than in the other ectodermal lineages (Figure 5A). To confirm these observations at a whole-genome scale, we analyzed the expression of 364 known direct YAP targets (Zanconato et al., 2015). The majority of these targets were significantly enriched in early epidermal lineage in comparison to other components of D4 neuruloid colonies (Figure 5B). These results confirm our initial observation in WT neuruloids that YAP is regulated and predominantly active in the epidermis lineage (Figures 2E, 3, and S3).

**Figure 5.**
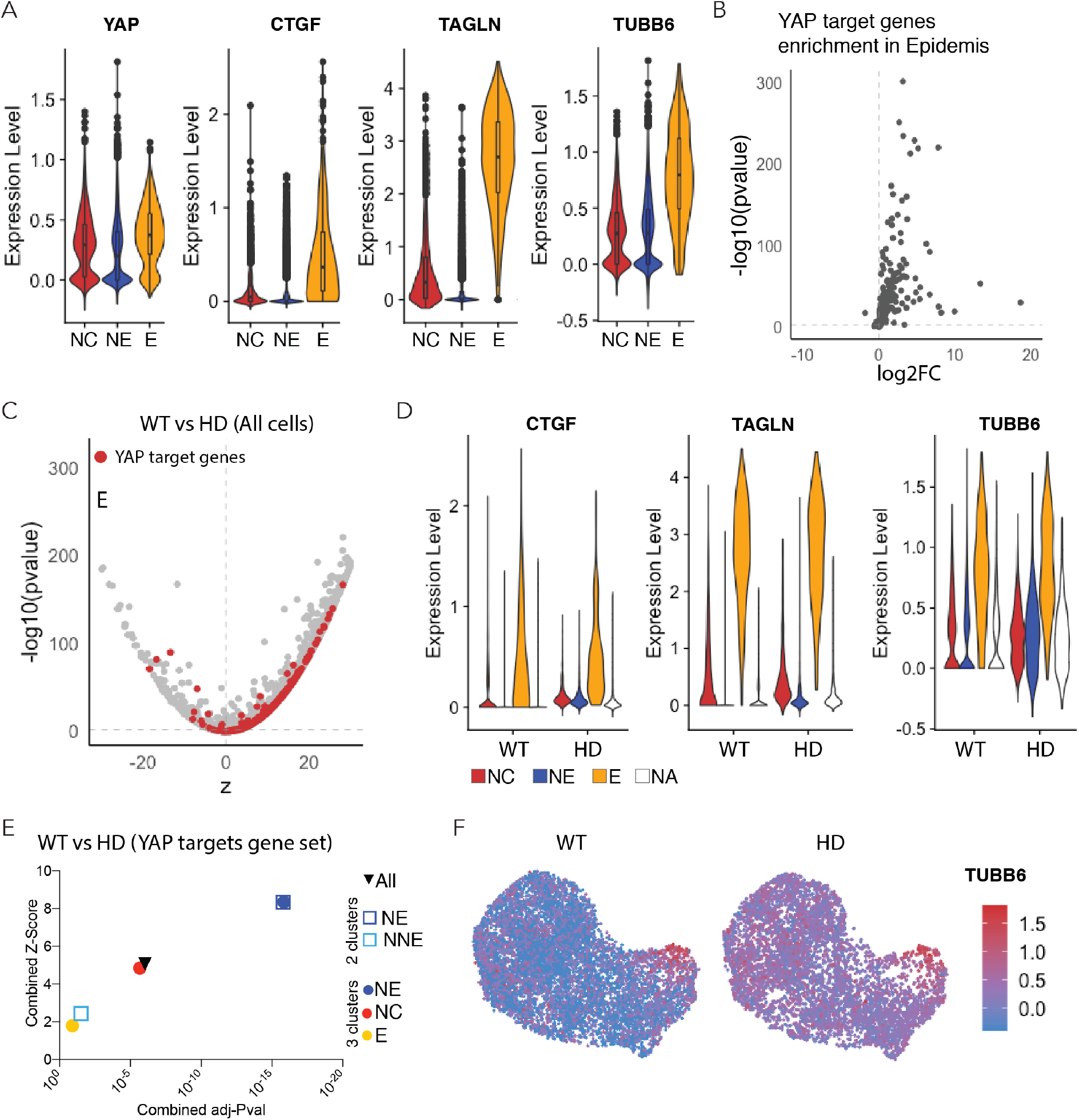
Expression of YAP target genes in WT and HD neuruloids. A) Violin plots show the expression of *YAP* and three representative target genes in the three clusters representing the main ectodermal lineages from scRNA-seq analysis of D4 neuruloids. Expression levels are normalized counts, calculated by Seurat for each gene and plotted on a logarithmic scale. B) A volcano plot shows the greater expression of several YAP target genes in epidermis compared to the remainder of the neuruloid. C) An analysis of differential gene expression from scRNA-seq data shows upregulation of YAP target genes (red) in HD with respect to WT neuruloids. D) Violin plots show the expression of three representative YAP target genes in the three lineage clusters from D4 WT and HD neuruloids. The color code is: neural crest (NC), red; neural ectoderm (NE), blue; epidermis (E), yellow; and unidentified (NA), white. The augmented expression of YAP target genes is more pronounced in the non-epidermal lineages. E) Gene-set enrichment analysis of YAP target genes confirms the effect. F) Analysis by uniform manifold approximation and projection (UMAP) shows the ectopic enhancement of a representative YAP target, *TUBB6*, in D4 WT and HD neuruloids.

We next used the scRNA-seq data to compare the molecular profiles of D4 early progenitors of neural ectoderm, neural crest, and epidermis in WT and HD neuruloids. Upon performing an analysis of differential gene expression, we ascertained that the majority of YAP target genes were significantly upregulated in HD compared to WT neuruloids (Figure 5C). This increased expression of YAP target genes was not observed in the epidermal lineage, for which YAP was highly active in both WT and HD neuruloids. Instead, YAP targets were ectopically activated in the NC and NE lineages of HD neuruloids by comparison to the WT counterparts (Figure 5D). To confirm this observation, we performed gene set-enrichment analyses on a gene set consisting of previously identified YAP target genes (Zanconato et al., 2015). In agreement with the analysis of differential gene expression, this approach demonstrated a significant increase in the expression of YAP targets in NC and especially NE lineages of D4 HD neuruloids (Figure 5E). At higher resolution, the analysis showed that YAP targets were significantly upregulated in the NE population of D4 HD neuruloids, whereas the upregulation in NNE was not significant (Figure 5E). Within the NNE lineage, only the NC cluster displayed a significant upregulation of YAP transcriptional activity. This phenomenon was well illustrated by the behavior of the YAP target gene TUBB6, which was normally expressed specifically in the early epidermal lineage, but became active throughout the neuruloid colony upon HD mutation (Figure 5F).

These results confirm that HD neuruloids display higher YAP activity during their formation. The data also reveal that such HD-associated activation of YAP does not directly affect the epidermal lineage, in which YAP is endogenously active, but primarily perturbs NE and NC, in which YAP becomes ectopically active as a result of the HD mutation.

### The HD phenotype in neuruloids results from ectopic YAP activation

In view of the increased YAP activity that we observed in HD neuruloids and the associated phenotypic consequences, we inquired whether inhibition of YAP transcriptional activity could rescue the HD phenotype. To block YAP activity, we used verteporfin, which prevents the interaction between YAP and TEADs and thereby inhibits the transcription of target genes (Liu-Chittenden et al., 2012). In agreement with the observation that a minimal level of YAP activity is required for ectodermal induction (Giraldez et al., 2021), administration of verteporfin during the initial three days of differentiation resulted in the loss of cell adhesion in both WT and HD samples. In conjunction with BMP4 induction, however, treatment with verteporfin from D3 was not toxic and did not affect the neuruloid phenotype of WT cells (Figures 6A,C and S8). This indicates that high level of YAP activity may be dispensable for epidermal specification.

**Figure 6.**
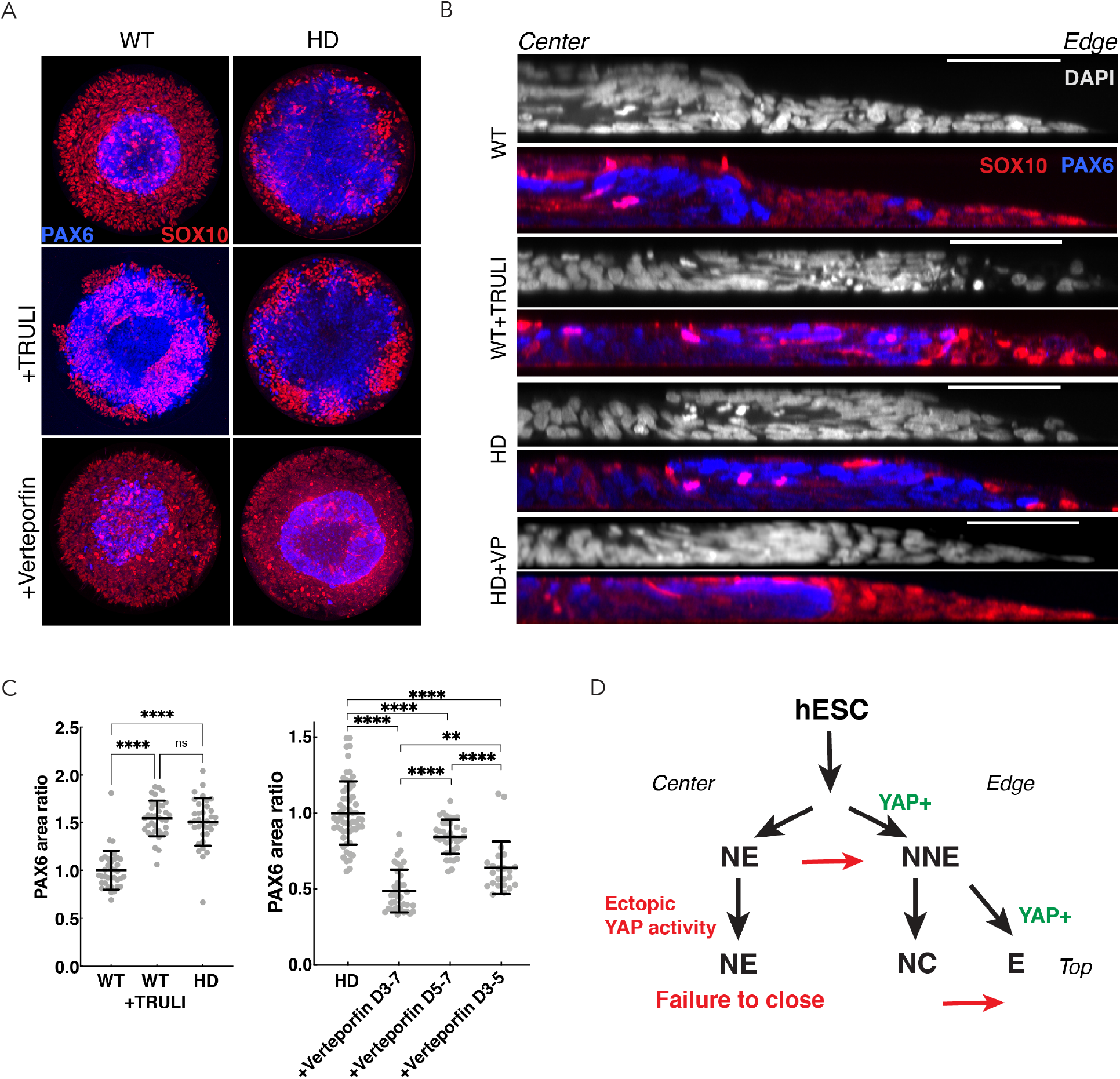
Perturbation of YAP activity in WT and HD neuruloids. A) Immunofluorescence images of D7 neuruloids show that the expansion of NE (PAX6, blue) at the expense of NC (PAX10, red) after exposure to 10 μM TRULI or in HD cells. Treatment with 0.3 μM verteporfin has no effect on WT colonies but partially rescues the effect of HD. B) Side views portray neuruloids under the conditions used in panel A. C) The ratios of PAX6+ areas to control values under various conditions confirm the similarity of TRULI-treated WT neuruloids to HD neuruloids and the suppressive effect of verteporfin. Rescue of this HD phenotype is stronger for early than for late exposure to verteporfin. ****, *p* < 0.0001 ; ns, *p* > 0.05 in unpaired *t*-tests comparing the different experimental conditions. D) In a model of neurulation, YAP activity (green) is posited to favor non-neuronal fates, and hyperactivity (red)—either from TRULI treatment or HD— yields characteristic abnormalities.

In D7 HD neuruloids, administration of verteporfin partially rescued aspects of the HD phenotype. Although the extended epidermal population observed in HD samples was maintained, the inhibition of YAP activity partially restored the NC (SOX10+) lineage and the condensation of the central NE (PAX6+) domain (Figures 6A-C and S8). This result accords with the observation that in D4 HD neuruloids, YAP target genes were significantly unregulated in the precursors of NC and NE, but not in the progenitors of the epidermal lineage (Figure 5).

Because YAP activity was elevated during the maturation of HD neuruloids from D4, we sought to determine the period during which the inhibition of YAP transcriptional activity was most important for rescuing the condensation of the central PAX6+ domain. We observed that verteporfin treatment rescued the HD-associated NE expansion if administrated together with BMP4 at D3, whereas it was unable to rescue the phenotype if added after D5 (Figure 6C).

These results indicate that the ectopic activity of YAP in HD neuruloids contributes to the malformation of their central PAX6 domains and that the effect is exerted early in neuruloid development.

## Discussion

In numerous organisms and a variety of tissues, the Hippo pathway plays important roles in development, homeostasis, and regeneration (Moya & Halder, 2019). Although exploring the role of this pathway during human development remains a challenge, organoid cultures derived from human embryonic stem cells offer simplified simulacra for investigation. Here we have identified YAP as an importantly determinant in neurulation, during which the protein’s activity is controlled—at least in part—by the Hippo pathway. Our data suggest that YAP plays a role in the specification of early ectodermal lineages, first in the choice between NNE and NE, and then that between epidermis and neural crest. Because YAP transcriptional activity is already robust at D4 in precursors of the epidermis, the Hippo pathway might play an early role in the specification of NNE during neurulation.

In line with previous reports (Yao et al., 2014; Huang et al., 2016), our results illustrate a reciprocal relationship between Hippo-YAP and BMP4-SAMD1/5 signaling during the development of the nervous system. In particular, YAP expression rises upon BMP4 stimulation, and YAP activation in turn enhances the cellular repose to BMP4 at various stages: initially by augmenting BMP4-dependent NNE induction, and later by favoring the specification of epidermis, which is known to occur at a high BMP4 concentration (Wilson and Hemmati-Brivanlou, 1995).

Noting the similarities between TRULI-treated neuruloids and those bearing HD mutations (Haremaki et al., 2019), we explored the potential contribution of YAP dysregulation to the HD phenotype. HD neuruloids demonstrated altered YAP regulation that resulted in ectopic activation of its transcriptional program, especially within the NE lineage. This HD-associated YAP hyperactivity might contribute directly to the failure of compaction in the central NE, even though this structure is devoid of YAP detectable by immunofluorescence.

Although individuals with Huntington’s Disease have not been reported to experience a failure in folding of the neural tube, there is evidence that some effects of disease-causing mutations occur during development but are activated at maturity (Barnat et al., 2020). For example, mice transiently exposed to the *Htt* mutant gene during development display an HD phenotype as adults (Molero et al., 2016; Wiatr et al., 2018). Owing to homeostatic and compensatory mechanisms, development *in vivo* is likely to be more robust than that in neuruloids. As a result, neuruloids offer a sensitive assay for subtle phenomena that might otherwise remain hidden. Because our data suggest that neurulation is perturbed in HD, exploring how the developing human embryo compensates for this challenge could yield insight into the dysfunction manifested at maturity.

Several studies support a role for YAP in the pathophysiology of HD. Postmortem cortical samples from HD patients display diminished YAP activity. Moreover, YAP activators ameliorate symptoms in HD mice and normalize the phenotypes of cultured cells (Mao et al., 2016; Mueller et al., 2018). In our data, YAP activation at D4 was elevated in HD neuruloids compared to WT controls, but at D7 YAP transcription was instead diminished (Figure S9). This reversal might reflect compensation for increased YAP activity so that development could proceed normally. The reversal might alternatively represent an end state of failed development. It will be interesting to investigate whether YAP hyperactivation during early neurulation leads to hypoactivity in mature neurons. Lats1/2 knockout mice demonstrate that Hippo signaling later regulates the number and differentiation of neural progenitor (Lavado et al., 2018), and postmortem analysis of HD patients shows significantly increased proliferation of subependymal neural progenitors (Curtis et al., 2005). These results suggest that the effects of *HTT* mutations on this process through YAP dysregulation bear on the pathophysiology of the disease.

Our study leaves open the question of the connection between Hippo signaling and HD. Huntingtin protein is known to play a role in vesicle trafficking and establishing apical-basal polarity, a prime regulator of Hippo signaling (Piccolo et al., 2014). The protein is also involved in the establishment and maintenance of intact epithelia, a function that is compromised by HD mutations (Galgoczi et al., 2021). Loss of intercellular adhesion could contribute directly to dysregulation of the Hippo pathway, reflected here as increased YAP activity. Lack of proper epithelial polarity could also result in the mislocalization of morphogen receptors and thus in increased sensitivity of BMP4/SMAD1/5 signaling, which in turn could affect the Hippo pathway. Future investigations of these mechanisms might reveal the molecular consequences of HD mutations and how mechanical cues interact with morphogenic signaling during embryonic development.

## Materials and Methods

### Cell culture

All hESC lines were grown in HUESM medium that was conditioned with mouse embryonic fibroblasts and supplemented with 20 ng/mL bFGF (MEF-CM; Deglincerti et al., 2016). Cells were grown on tissue culture dishes coated with Geltrex solution (Life Technologies) and tested for mycoplasma at two-month intervals.

### Generation of YAP-GFP hESC lines

Carboxy-terminal tagging of endogenous YAP alleles with GFP was obtained by using a donor plasmids, which was kindly donated by the Liphardt laboratory (Franklin et al., 2020). This contains an upstream homology arm approximately 1 kB in length; a cassette encoding GFP, a P2A site, puromycin resistance, and a stop codon; and a downstream homology arm also about 1 kB long. RUES2 (WT) and isogenic 56CAG (HD) hESC lines were co-transfected with Px330-Cas9-sgYAP (sgRNA sequence targeting YAP locus: TTAGAATTCAGTCTGCCTGA), donor YAP-eGFP-P2A-puromycin resistance, ePB-H2B-mCherry-BSDa, and transposase. Nucleofection was performed with program B-016 on an Amaxa Nucleofector II (Lonza). Cells were then grown for ten days under selection by 1 μg/mL puromycin and 10 μg/mL blasticidin. Multiple clones from each line were selected for characterization by PCR genotyping with the following oligonucleotides:

GFP integration:

YAP to exon9-1F: CAGGGGTAATTACGGAAGCA
mEGFP_R: CTGAACTTGTGGCCGTTTAC

Heterozygotic versus homozygotic:

Yap Het/Homo check LHA F: GGTGATACTATCAACCAAAGCACCC
YAP Het/Homo check R: CATCCATCATCCAAACAGGCTCAC

### Neuruloid culture

To generate neuruloids (Haremaki et al., 2019), we coaated micropatterned glass cover slips (CYTOO Arena A, Arena 500 A, Arena EMB A) for 3 hr at 37 °C with 10 μg/mL recombinant laminin-521 (BioLamina, LN521-05) diluted in PBS+/+ (Gibco). Single-cell suspensions were then incubated for 3 hr at 37 °C in HUESM medium supplemented with 20 ng/mL bFGF and 10 μM Rock inhibitor Y27632. The micropattern culture was then washed once with PBS+/+ and incubated with HUESM with 10 μM SB431542 and 0.2 μM LDN 193189. On D3, the medium was replaced with HUESM containing 10 μM SB431542 and 3 ng/mL BMP4. On D5 the medium was replaced with the same fresh medium then incubated until D7.

### Neural crest and epidermis differentiation

Direct differentiation of neural crest was performed by a published method (Tchieu et al., 2017). RUES2 hESCs were cultured for three days in E6 medium (STEMCELL, 05946) containing 1 ng/mL BMP4, 10 μM SB431542, and 600 nM CHIR, then transferred for four days to 10 μM SB431542 and 1.5 μM CHIR.

Induction of epidermis was conducted on Transwell (Corning, CLS3413). RUES2 hESCs were cultured for three days in E6 medium (STEMCELL, 05946) containing 10 μM SB431542 (top of the transwell) and 10 μM SB431542 and 10 ng/mL BMP4 (bottom of the transwell). After the media had been replaced with 10 μM SB431542 and 0.2 μM LDN193189 (top and bottom), the cells were incubated for an additional four days.

### Immunofluorescence

Micropatterned coverslips were fixed with 4 % paraformaldehyde (Electron Microscopy Sciences 15713) in warm medium for 30 min, rinsed three times with PBS−/−, and blocked and permeabilized for 30 min with 3 % normal donkey serum (Jackson Immunoresearch 017-000-121) and 0.5 % Triton X-100 (Sigma 93443) in PBS−/−. Specimens were incubated with primary antibodies for 1.5 hr at room temperature, washed three times for 5 min each in PBS−/−, incubated with secondary antibodies conjugated with Alexa 488, Alexa 555, Alexa 594 or Alexa 647 (Molecular Probes) at 1/1000 dilution. After 30 min incubation with 100 ng/mL 4’,6-diamidino-2-phenylindole (DAPI; Thermo Fisher Scientific D1306), specimens were washed three times with PBS−/−. Coverslips were mounted on slides using ProLong Gold antifade mounting medium (Molecular Probes P36934).

Antibodies used:

YAP (Santa Cruz Biotechnology 101199)
TFAP2A (DSHB 3B5 concentrate)
PAX6 (BD Biosciences 561462)
SOX10 (R&D Systems AF2864)
KRT18 (Abcam ab194130)

### Imaging and analysis

Confocal images were acquired on a Zeiss Inverted LSM 780 laser-scanning confocal microscope with a 10X, 20X, or 40X water-immersion objective. YAP-GFP Live reporter imaging was performed with a Zeiss AxioObserver z1 or spinning-disk microscope (Cell Voyager CV1000, Yokogawa).

For the acquisition of radial profiles, images were preprocessed to display the maximum-intensity projection (MIP) of at least four z-stacks. MIP images were subsequently imported into Python (czifile 2019.7.2). Individual colonies were cropped and the radial intensity profile of fluorescence was calculated for each image starting from the center. Results were reported in arbitrary fluorescence units.

For quantitative analysis of single cells (Figure 2D), individual nuclei were segmented in the DAPI channel in each z-slice of a confocal image by a published procedure (Etoc et al., 2016). Twelve images representative of the data were selected to train Ilastik, an interactive learning and segmentation toolkit, to classify each pixel into two categories: nucleus or background. The pixels in the nucleus class were then used to create a binary mask. The original z-slice image was then processed to detect seeds based on a sphere filter with a manually set threshold. Watershedding was then applied from the defined seeds into the mask of the nuclei as defined previously by Ilastik, resulting in a set of segmented nuclei. The median intensity of PAX6, SOX10, or YAP immunofluorescence was then measured for each segmented nucleus.

### Single-cell RNA sequencing

Micropatterned glass coverslips with neuruloids 500 μm in diameter were grown in 3 ng/mL BMP4 from RUES2 hESCs and the isogenic 56CAG HD line. On D4, approximately half of the neuruloids on each coverslip were scraped off and treated for 10 min at 37 °C with Accutase (Stemcell Technologies). The remaining neuruloids were fixed for immunofluorescence analysis as a quality control. After dissociation, the cells were washed three times in PBS−/− (Gibco) with 0.04 % BSA and strained through a 40 μm tip strainer (Flowmi 136800040). The number and viability of cells were determined by exclusion of trypan blue on a Countess II automated cell counter. The samples were separately loaded for capture with the Chromium System using Single Cell 3’ v2 reagents (10× Genomics). Following cell capture and lysis, cDNA was synthesized and amplified according to the manufacturer’s instructions. The resulting libraries were sequenced on NovaSeq 6000 SP flowcell. The Cell Ranger software pipeline (v 2.0.2) was used to create FASTQ files that aligned to the hg19 genome with default parameters. Filtered gene-expression matrices of genes *versus* detected barcodes (cells) with counts of unique molecular identifiers (UMI) were generated and used for subsequent analyses. These data are available through NCBI GEOXXXX accession

### Single-cell RNA analysis

The sequencing data were analyzed using Cellranger count (version 3.0.2) and aggregated with the Aggr function and default settings. The aggregated datasets were processed with Seurat (version 3.2.3) (Stuart et al., 2019). Quality control and filtering were performed with Scater Bioconductor (version 1.16.1) to identify and remove poor-quality cells characterized by low library sizes, low numbers of expressed genes, and low or high numbers of mitochondrial reads (McCarthy et al., 2017). Following a cut-off step (counts > 6000; genes > 2600; 3 % < mitochondrial genes < 11 %), we obtained about 8000 cells in WT and roughly 6000 cells for HD. Counts of genes in each cell were normalized and log_10_-transformed. In order to merge the datasets using Seurat’s FindIntegrationAnchors and IntegrateData methods, we identified integration anchors for the WT and HD datasets by means of the top 2000 variable genes and first 20 principal components. The cell cycle phase was estimated by Seurat with default setting and data were rescaled to remove the effects of cell cycle and mitochondrial transcript expression. Clustering of cell was performed with Seurat’s findClusters function by use of the first 20 principal components and with two resolutions—0.5 and 0.1—to yield sets of seven clusters and two clusters, respectively. The seven-cluster set was further aggregated by manual curation of cell-type specific markers into four clusters. UMAP dimension reduction implemented within Seurat was applied to the normalized, integrated data and the cluster sets projected and visualized on the resulting UMAP coordinates. Visualization of normalized expression values on UMAP coordinates and as violin plots were performed using Seurat’s FeaturePlot and VlnPlot, respectively. Differential gene expression and analysis between clusters and samples were performed by MAST (version 1.8.2; Finak et al., 2015). YAP targets were retrieved from msigDB C6 gene sets and enrichment with differential expression tests assessed by single-cell GSEA implemented within MAST.

## Acknowledgments

FMP, TH, QT, TLL and AHB were supported by the CHDI foundation (A-9423). NRK was supported by NIGMS grant T32GM007739. RDS was supported by EMBO-LTF-254-2019. KG was supported by NIDCD grant R21DC016984. AJH is an Investigator of Howard Hughes Medical Institute.

## Author contributions

FMP designed the experiments, analyzed the results, generated the figures and wrote the manuscript. NRK provided a reagent and wrote the manuscript. TH, QT and TLL executed the experiments. RDS provided analytical tools. TSC and JDL performed bioinformatics analysis. FE contributed to initial conceptualization and provided analytical tools. KG provided a reagent and edited the manuscript. AJH edited the manuscript and figures and supervised the study. AHB supervised the study.

## References

Barnat, M., Capizzi, M., Aparicio, E., Boluda, S., Wennagel, D., Kacher, R., Kassem, R., Lenoir, S., Agasse, F., Braz, B. Y., Liu, J.-P., Ighil, J., Tessier, A., Zeitlin, S. O., Duyckaerts, C., Dommergues, M., Durr, A., & Humbert, S. (2020). Huntington’s disease alters human neurodevelopment. Science, 369(6505), 787–793. https://doi.org/10.1126/science.aax3338

Curtis, M. A., Penney, E. B., Pearson, J., Dragunow, M., Connor, B., & Faull, R. L. M. (2005). The distribution of progenitor cells in the subependymal layer of the lateral ventricle in the normal and Huntington’s disease human brain. Neuroscience, 132(3), 777–788. https://doi.org/10.1016/j.neuroscience.2004.12.051

Etoc, F., Metzger, J., Ruzo, A., Kirst, C., Yoney, A., Ozair, M. Z., Brivanlou, A. H., & Siggia, E. D. (2016). A Balance between Secreted Inhibitors and Edge Sensing Controls Gastruloid Self-Organization. Developmental Cell, 39(3), 302–315. https://doi.org/10.1016/j.devcel.2016.09.016

Finak, G., McDavid, A., Yajima, M., Deng, J., Gersuk, V., Shalek, A. K., Slichter, C. K., Miller, H. W., McElrath, M. J., Prlic, M., Linsley, P. S., & Gottardo, R. (2015). MAST: A flexible statistical framework for assessing transcriptional changes and characterizing heterogeneity in single-cell RNA sequencing data. Genome Biology, 16(1), 278. https://doi.org/10.1186/s13059-015-0844-5

Franklin, J. M., Ghosh, R. P., Shi, Q., Reddick, M. P., & Liphardt, J. T. (2020). Concerted localization-resets precede YAP-dependent transcription. Nature Communications, 11(1), 4581. https://doi.org/10.1038/s41467-020-18368-x

Galgoczi, S., Ruzo, A., Markopoulos, C., Yoney, A., Phan-Everson, T., Haremaki, T., Metzger, J. J., Etoc, F., & Brivanlou, A. H. (2021). *Huntingtin CAG expansion impairs germ layer patterning in synthetic human gastruloids through polarity defects* [Preprint]. Developmental Biology. https://doi.org/10.1101/2021.02.06.430005

Giraldez, S., Stronati, E., Huang, L., Hsu, H.-T., Abraham, E., Jones, K. A., & Estaras, C. (2021). *YAP1 Regulates the Self-organized Fate Patterning of hESCs-Derived Gastruloids* [Preprint]. Developmental Biology. https://doi.org/10.1101/2021.03.12.434631

Haremaki, T., Metzger, J. J., Rito, T., Ozair, M. Z., Etoc, F., & Brivanlou, A. H. (2019). Self-organizing neuruloids model developmental aspects of Huntington’s disease in the ectodermal compartment. Nature Biotechnology, 37(10), 1198–1208. https://doi.org/10.1038/s41587-019-0237-5

Huang, Z., Hu, J., Pan, J., Wang, Y., Hu, G., Zhou, J., Mei, L., & Xiong, W.-C. (2016). YAP stabilizes SMAD1 and promotes BMP2-induced neocortical astrocytic differentiation. Development, 143(13), 2398–2409. https://doi.org/10.1242/dev.130658

Kastan, N., Gnedeva, K., Alisch, T., Petelski, A. A., Huggins, D. J., Chiaravalli, J., Aharanov, A., Shakked, A., Tzahor, E., Nagiel, A., Segil, N., & Hudspeth, A. J. (2021). Small-molecule inhibition of Lats kinases may promote Yap-dependent proliferation in postmitotic mammalian tissues. Nature Communications, 12(1), 3100. https://doi.org/10.1038/s41467-021-23395-3

Lavado, A., Park, J. Y., Paré, J., Finkelstein, D., Pan, H., Xu, B., Fan, Y., Kumar, R. P., Neale, G., Kwak, Y. D., McKinnon, P. J., Johnson, R. L., & Cao, X. (2018). The Hippo Pathway Prevents YAP/TAZ-Driven Hypertranscription and Controls Neural Progenitor Number. Developmental Cell, 47(5), 576–591.e8. https://doi.org/10.1016/j.devcel.2018.09.021

Liu-Chittenden, Y., Huang, B., Shim, J. S., Chen, Q., Lee, S.-J., Anders, R. A., Liu, J. O., & Pan, D. (2012). Genetic and pharmacological disruption of the TEAD-YAP complex suppresses the oncogenic activity of YAP. Genes & Development, 26(12), 1300–1305. https://doi.org/10.1101/gad.192856.112

Mao, Y., Chen, X., Xu, M., Fujita, K., Motoki, K., Sasabe, T., Homma, H., Murata, M., Tagawa, K., Tamura, T., Kaye, J., Finkbeiner, S., Blandino, G., Sudol, M., & Okazawa, H. (2016). Targeting TEAD/YAP-transcription-dependent necrosis, TRIAD, ameliorates Huntington’s disease pathology. Human Molecular Genetics, ddw303. https://doi.org/10.1093/hmg/ddw303

McCarthy, D. J., Campbell, K. R., Lun, A. T. L., & Wills, Q. F. (2017). Scater: Pre-processing, quality control, normalization and visualization of single-cell RNA-seq data in R. Bioinformatics, btw777. https://doi.org/10.1093/bioinformatics/btw777

Molero, A. E., Arteaga-Bracho, E. E., Chen, C. H., Gulinello, M., Winchester, M. L., Pichamoorthy, N., Gokhan, S., Khodakhah, K., & Mehler, M. F. (2016). Selective expression of mutant huntingtin during development recapitulates characteristic features of Huntington’s disease. Proceedings of the National Academy of Sciences, 113(20), 5736–5741. https://doi.org/10.1073/pnas.1603871113

Moya, I. M., & Halder, G. (2019). Hippo–YAP/TAZ signalling in organ regeneration and regenerative medicine. Nature Reviews Molecular Cell Biology, 20(4), 211–226. https://doi.org/10.1038/s41580-018-0086-y

Mueller, K. A., Glajch, K. E., Huizenga, M. N., Wilson, R. A., Granucci, E. J., Dios, A. M., Tousley, A. R., Iuliano, M., Weisman, E., LaQuaglia, M. J., DiFiglia, M., Kegel-Gleason, K., Vakili, K., & Sadri-Vakili, G. (2018). Hippo Signaling Pathway Dysregulation in Human Huntington’s Disease Brain and Neuronal Stem Cells. Scientific Reports, 8(1), 11355. https://doi.org/10.1038/s41598-018-29319-4

Piccolo, S., Dupont, S., & Cordenonsi, M. (2014). The Biology of YAP/TAZ: Hippo Signaling and Beyond. Physiological Reviews, 94(4), 1287–1312. https://doi.org/10.1152/physrev.00005.2014

Stuart, T., Butler, A., Hoffman, P., Hafemeister, C., Papalexi, E., Mauck, W. M., Hao, Y., Stoeckius, M., Smibert, P., & Satija, R. (2019). Comprehensive Integration of Single-Cell Data. Cell, 177(7), 1888–1902.e21. https://doi.org/10.1016/j.cell.2019.05.031

Tchieu, J., Zimmer, B., Fattahi, F., Amin, S., Zeltner, N., Chen, S., & Studer, L. (2017). A Modular Platform for Differentiation of Human PSCs into All Major Ectodermal Lineages. Cell Stem Cell, 21(3), 399–410.e7. https://doi.org/10.1016/j.stem.2017.08.015

Wiatr, K., Szlachcic, W. J., Trzeciak, M., Figlerowicz, M., & Figiel, M. (2018). Huntington Disease as a Neurodevelopmental Disorder and Early Signs of the Disease in Stem Cells. Molecular Neurobiology, 55(4), 3351–3371. https://doi.org/10.1007/s12035-017-0477-7

Wilson, P. A., & Hemmati-Brivanlou, A. (1995). Induction of epidermis and inhibition of neural fate by Bmp-4. Nature, 376(6538), 331–333. https://doi.org/10.1038/376331a0

Wu, Z., & Guan, K.-L. (2021). Hippo Signaling in Embryogenesis and Development. Trends in Biochemical Sciences, 46(1), 51–63. https://doi.org/10.1016/j.tibs.2020.08.008

Yao, M., Wang, Y., Zhang, P., Chen, H., Xu, Z., Jiao, J., & Yuan, Z. (2014). BMP2-SMAD Signaling Represses the Proliferation of Embryonic Neural Stem Cells through YAP. Journal of Neuroscience, 34(36), 12039–12048. https://doi.org/10.1523/JNEUROSCI.0486-14.2014

Zanconato, F., Forcato, M., Battilana, G., Azzolin, L., Quaranta, E., Bodega, B., Rosato, A., Bicciato, S., Cordenonsi, M., & Piccolo, S. (2015). Genome-wide association between YAP/TAZ/TEAD and AP-1 at enhancers drives oncogenic growth. Nature Cell Biology, 17(9), 1218–1227. https://doi.org/10.1038/ncb3216

