## Supplemental Figures for "Role of YAP in early ectodermal specification and a Huntington’s Disease model of human neurulation"

**Supplemental figures and legends**

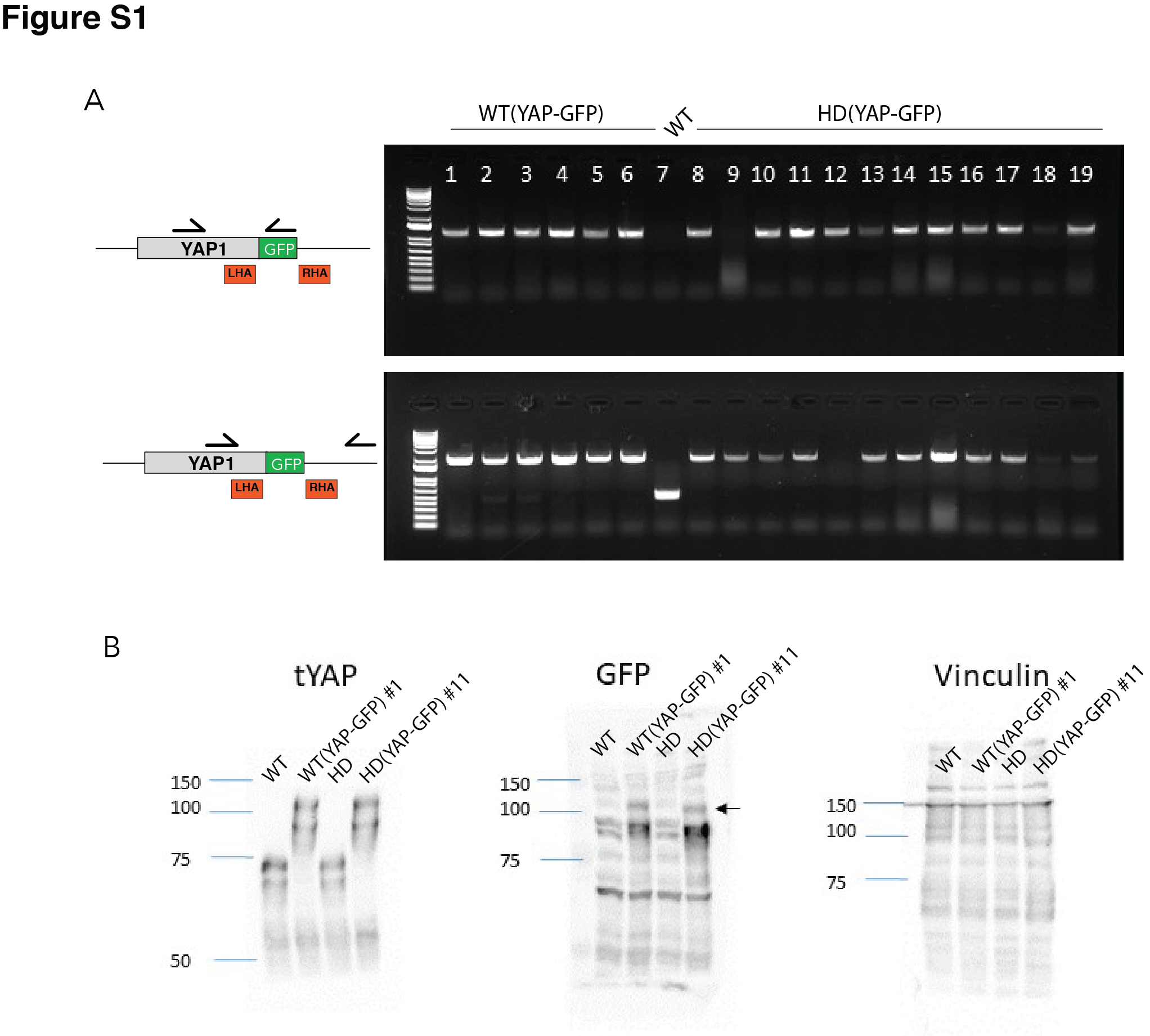

**Figure S1. Screening of WT and HD YAP-GFP reporter lines.** A) Genotypic screening of hESC clones from WT and HD backgrounds with PCR oligonucleotides (arrows) shows that multiple colonies underwent biallelic insertion of the GFP sequence. B) Immunoblot for total YAP (tYAP) and GFP from one clone of WT (YAP‑GFP) and one from HD (YAP‑GFP) confirmed the expression of YAP‑GFP fusion proteins. Untransfected WT and HD lines provide negative controls; vinculin provides a loading control.

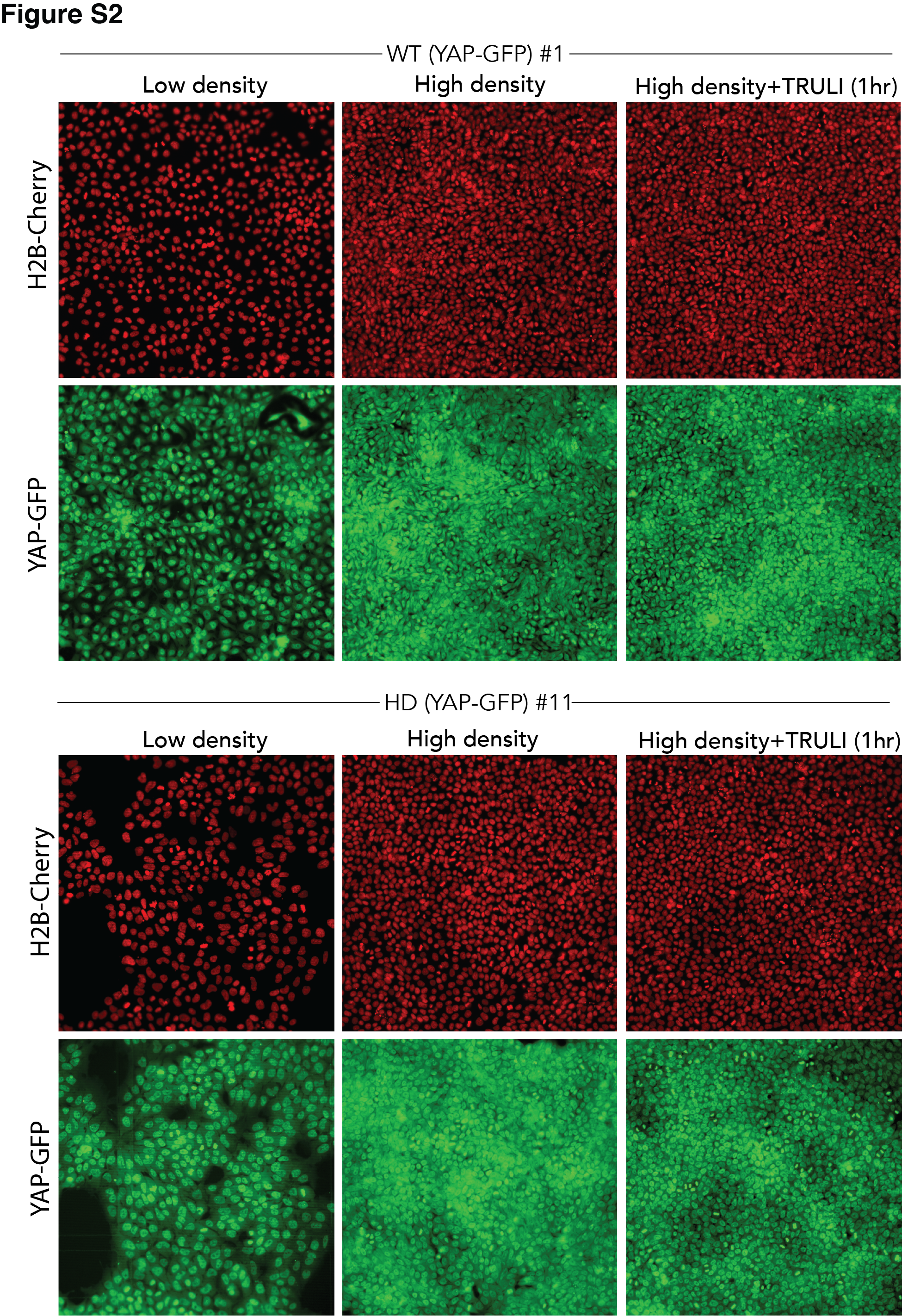

**Figure S2. Validation of WT and HD YAP-GFP reporter lines.** Fluorescent images of living WT and HD hESC clones expressing endogenous YAP‑GFP and H2b‑Cherry show the effect on the nuclear-cytoplasmic flux of YAP of cellular density and of treatment with 10 μM TRULI for 1 hr.

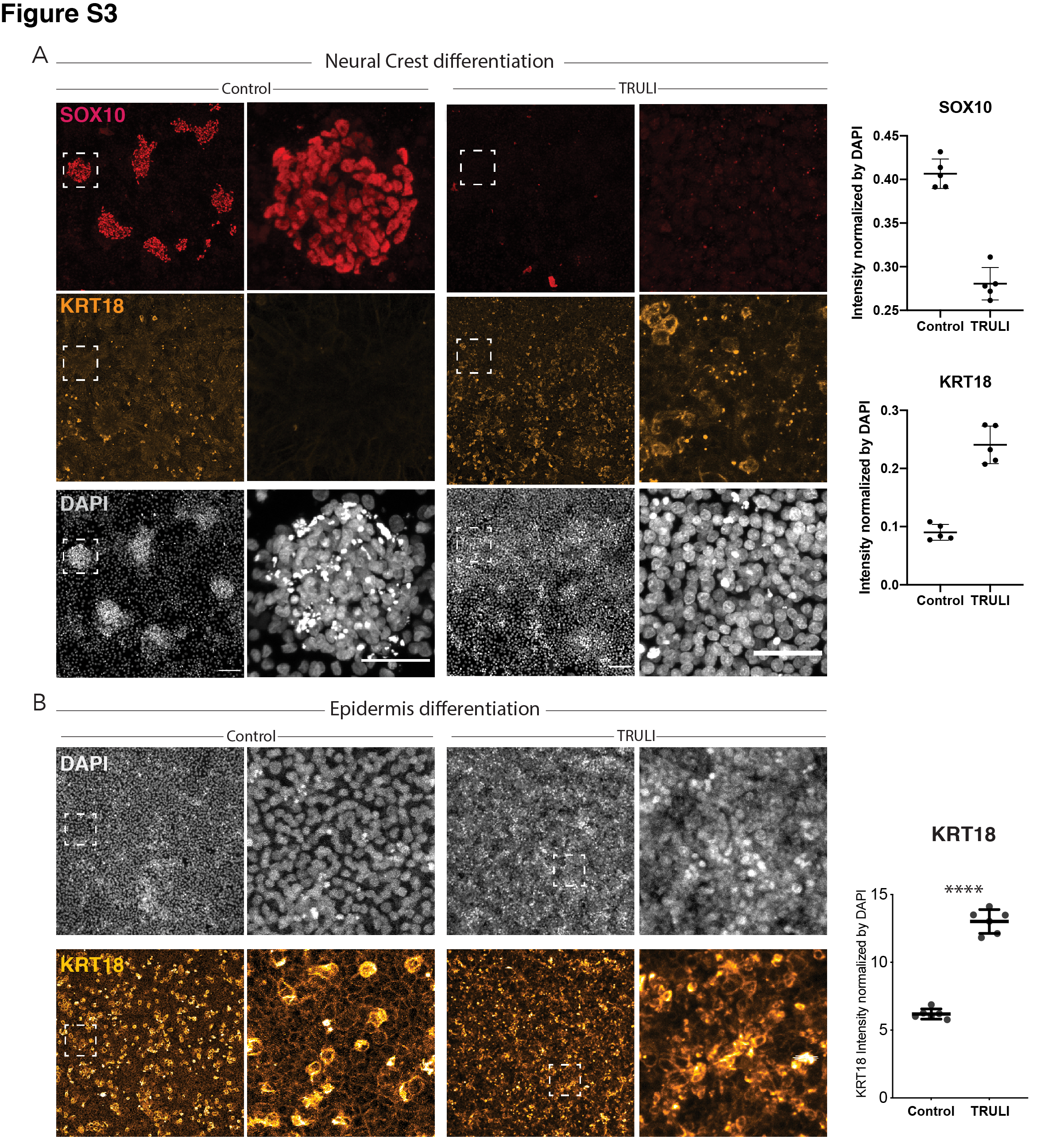

**Figure S3. Effect of sustained YAP activation on differentiation of neural crest and epidermis.** A) In immunofluorescence images of hESC after 10 days of a differentiation protocol for neural crest without (control) or in the presence of 10 μM TRULI treatment precludes the formation of NC (SOX10, red) and yields significantly more epidermal cells (KRT18, yellow). At the right, plots quantify the effect of TRULI treatment in this protocol on SOX10 and KRT18 signals, which are normalized to the DAPI signal. B) Immunofluorescence images of hESC after 10 days of a differentiation protocol for epidermis without (control) or in the presence of 10 μM TRULI reveal that TRULI treatment increases the intensity of KRT18 labeling. At the right, plots quantify the change in the KRT18 signal normalized to the DAPI signal and confirm that YAP activation results in the expression of KRT18.

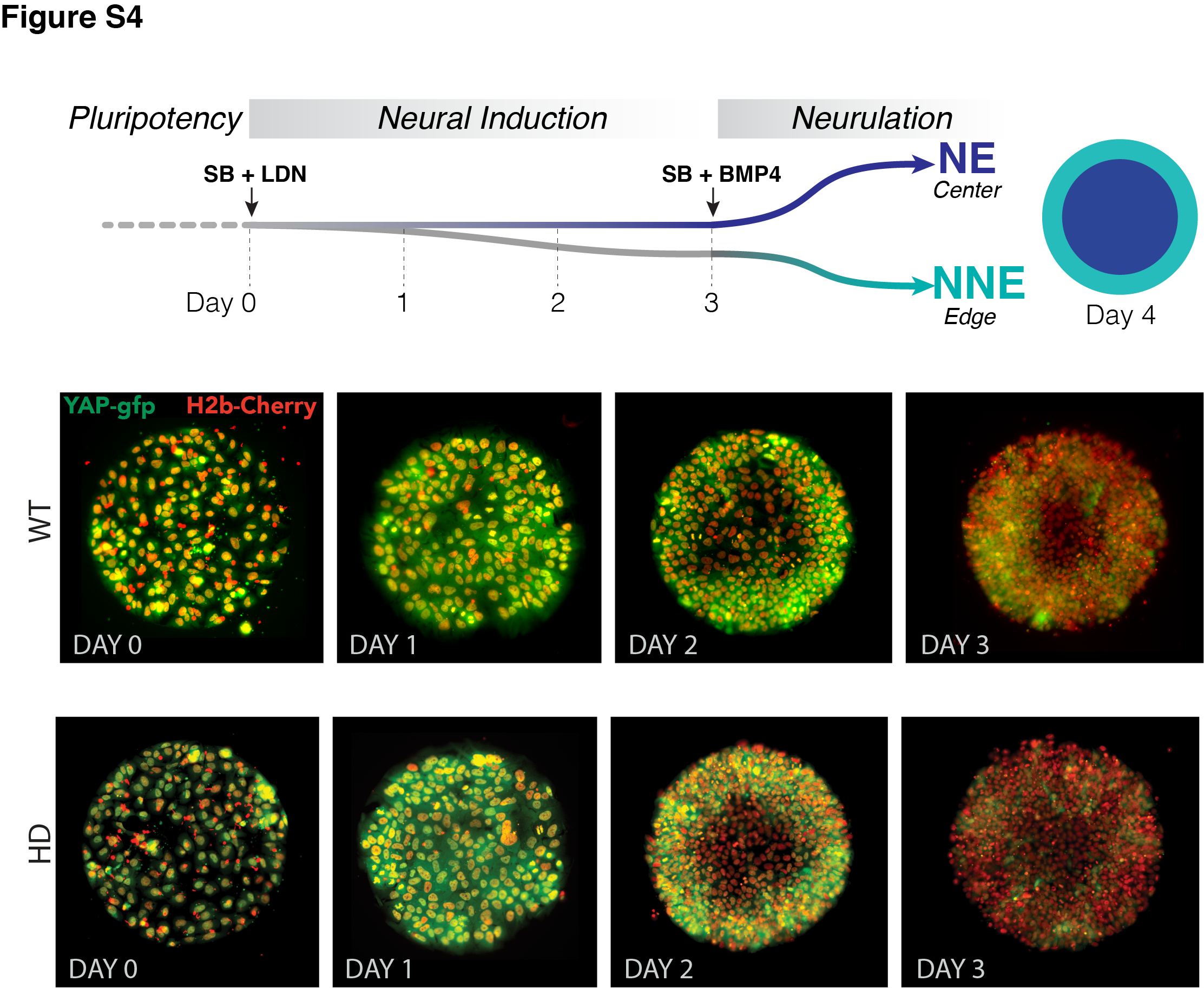

**Figure S4. Dynamics of YAP localization in early WT and HD neuruloids.** A) During the first four days of the differentiation protocol, pluripotent hESCs seeded on a micropattern undergo three days of neural induction (SB+LDN) followed by BMP4 induction of neurulation. B) Fluorescent images of living YAP‑GFP and H2b‑Cherry hESC colonies during neural induction show the progressive shift of YAP from nuclei into cytoplasm and a reduction in expression level for both WT and HD samples.

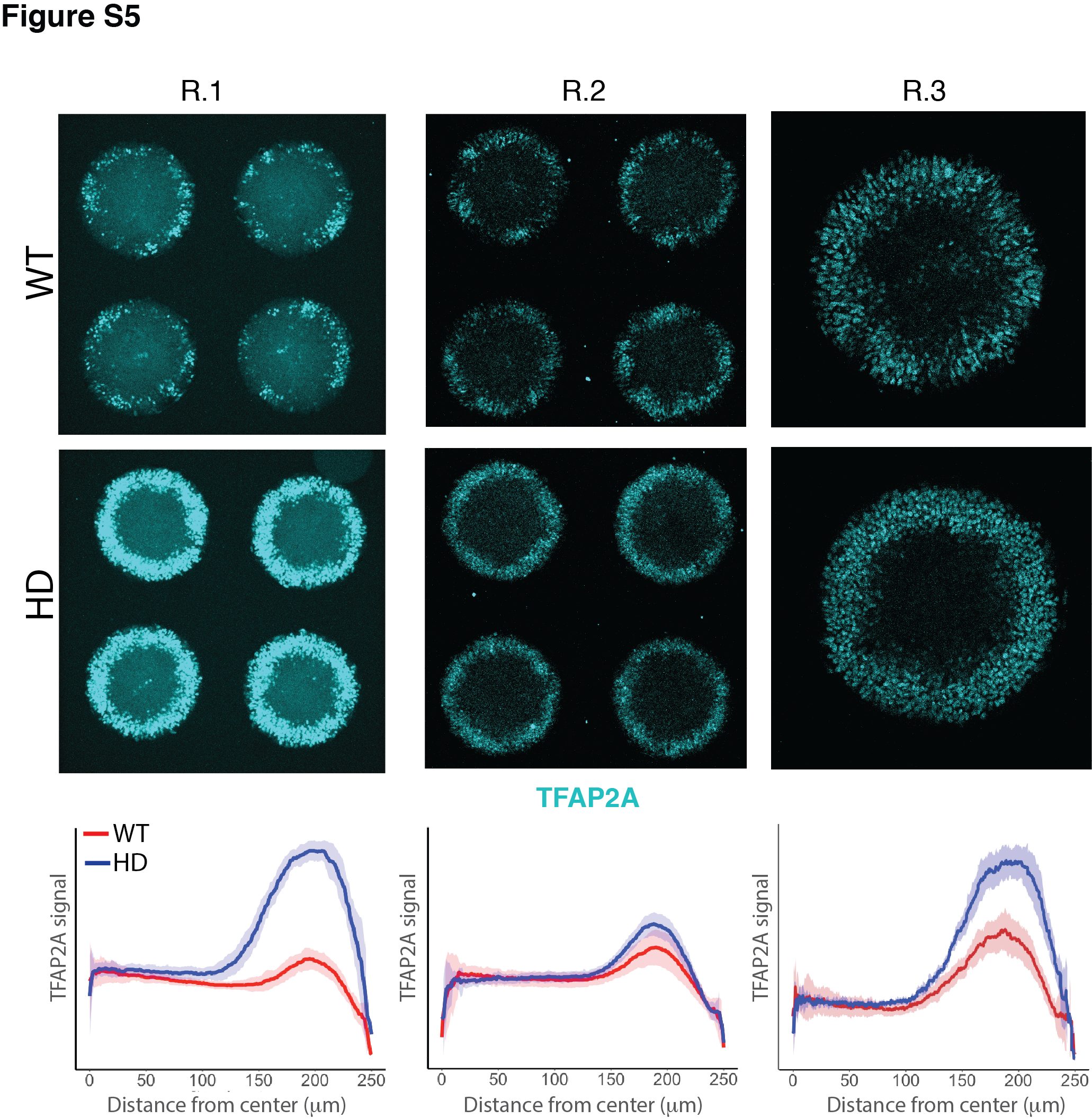

**Figure S5. Induction of NNE in WT and HD D4 neuruloids.** Immunofluorescence images of D4 neuruloids demonstrate enhanced expression of TFAP2A at the perimeters of HD colonies in three independent experiments. At the bottom, the radial distribution of the nuclear TFAP2A signal confirms this observation in multiple colonies (R.1, *n* = 16; R.2, *n* = 9; R.3, *n* = 12).

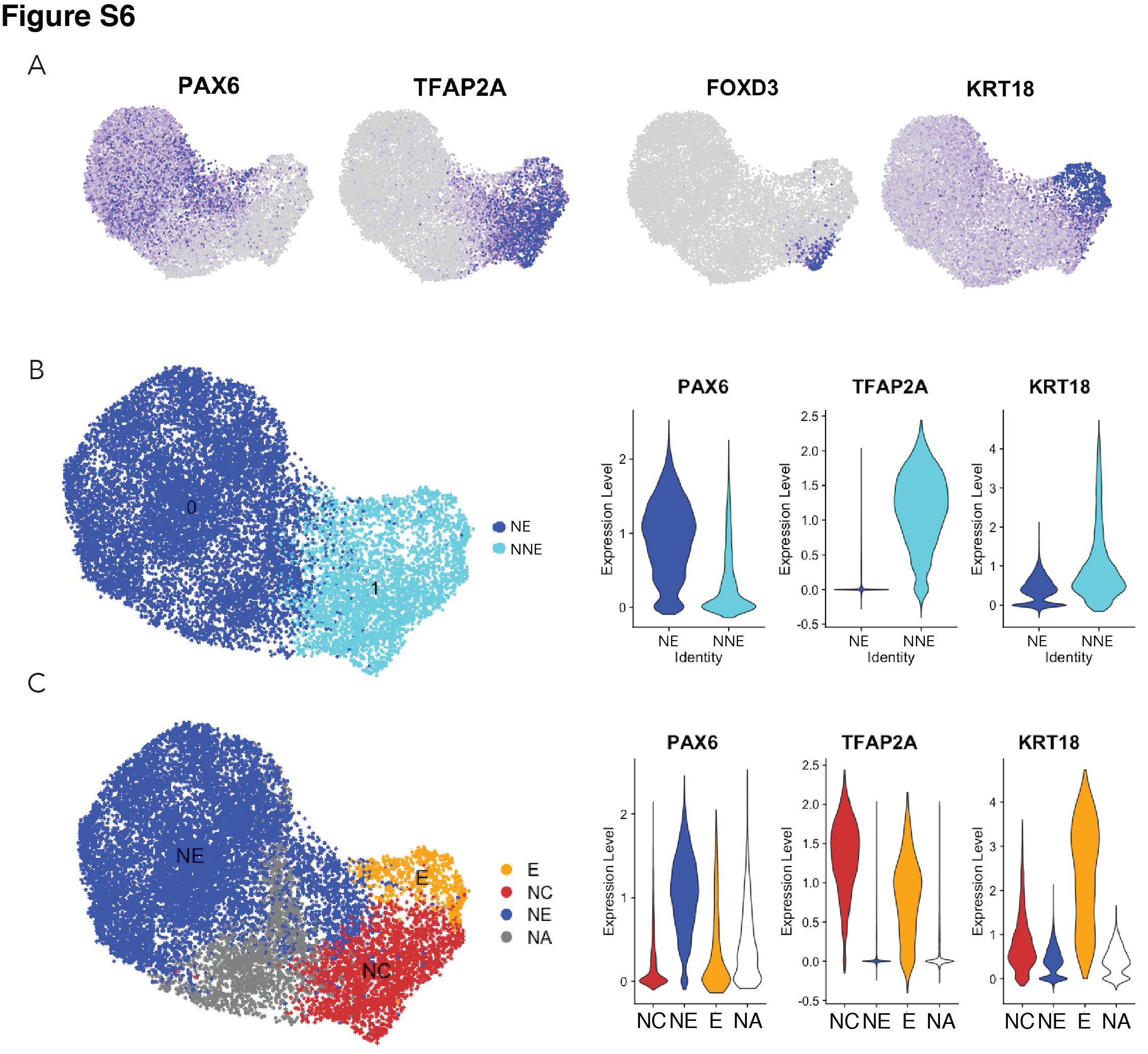

**Figure S6. Ectodermal lineage annotation in scRNA‑seq from D4 neuruloids.** A) Uniform manifold approximation and projection (UMAP) shows expression of key ectodermal lineage makers: *PAX6* identifies neural ectoderm (NE) and *TFAP2A* delineates non-neural ectoderm (NNE). Within the NNE, *FOXD3* identifies progenitors of neural crest (NC) and *KRT18* marks early epidermis (E). B) On the left, a plot documents the separation of NE and NNE in UMAP. At the right, violin plots show the expression of lineage genes in the NE and NNE. C) On the left, a plot demonstrates the separation of the three ectodermal linages in UMAP. NA represent the population of unidentified cells. At the right, violin plots show the expression of lineage genes in the three populations.

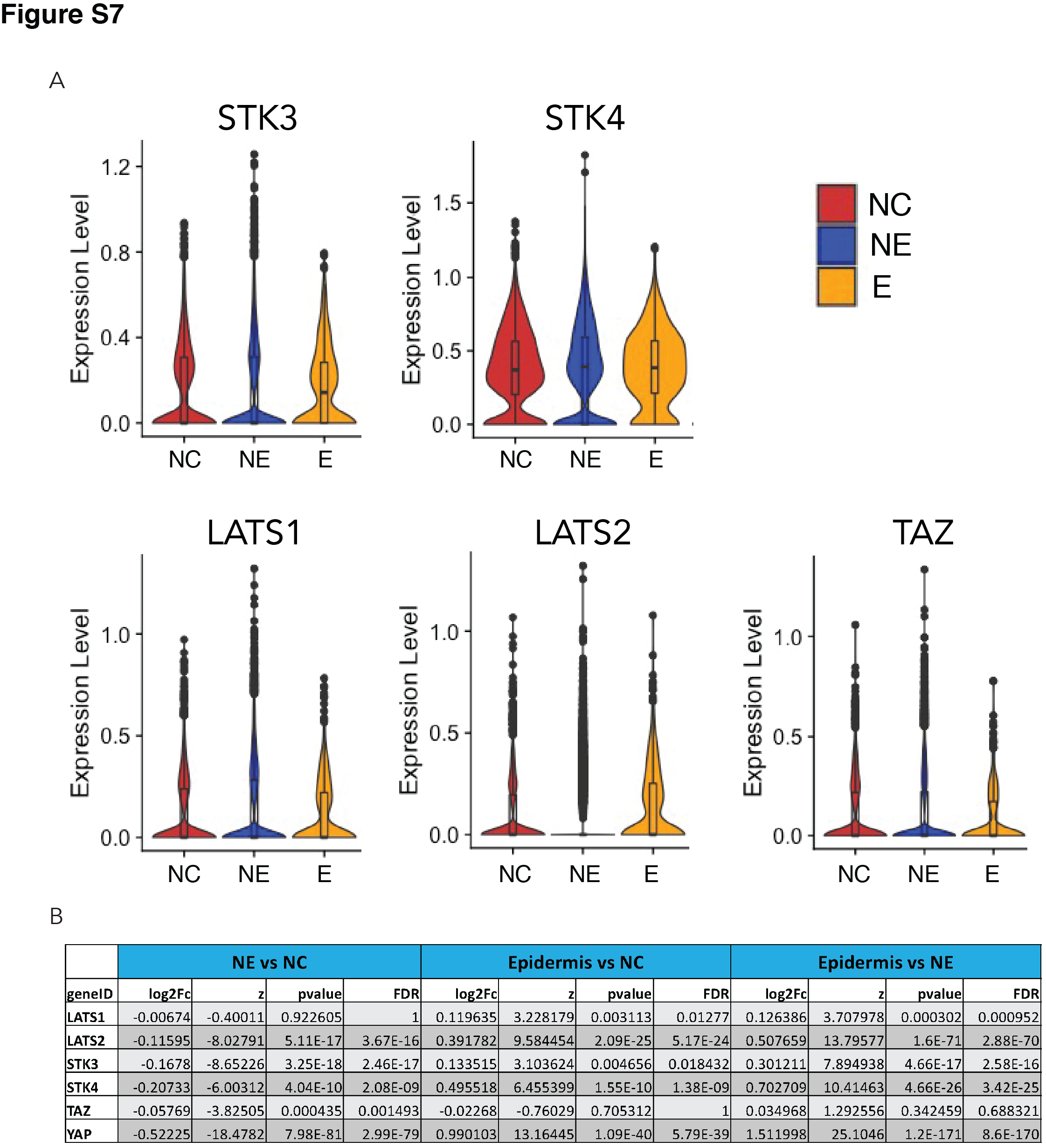

**Figure S7. Expression of key component of the Hippo pathway in D4 neuruloids.** A) Violin plots document the expression of components of the Hippo pathway in the three ectodermal lineages. B) A table summarizes the statistics for the pairwise differential-expression analysis of components of the Hippo pathway. *YAP* and key Hippo-pathway components can be found in all three lineages, but are more strongly expressed in NC and epidermis.

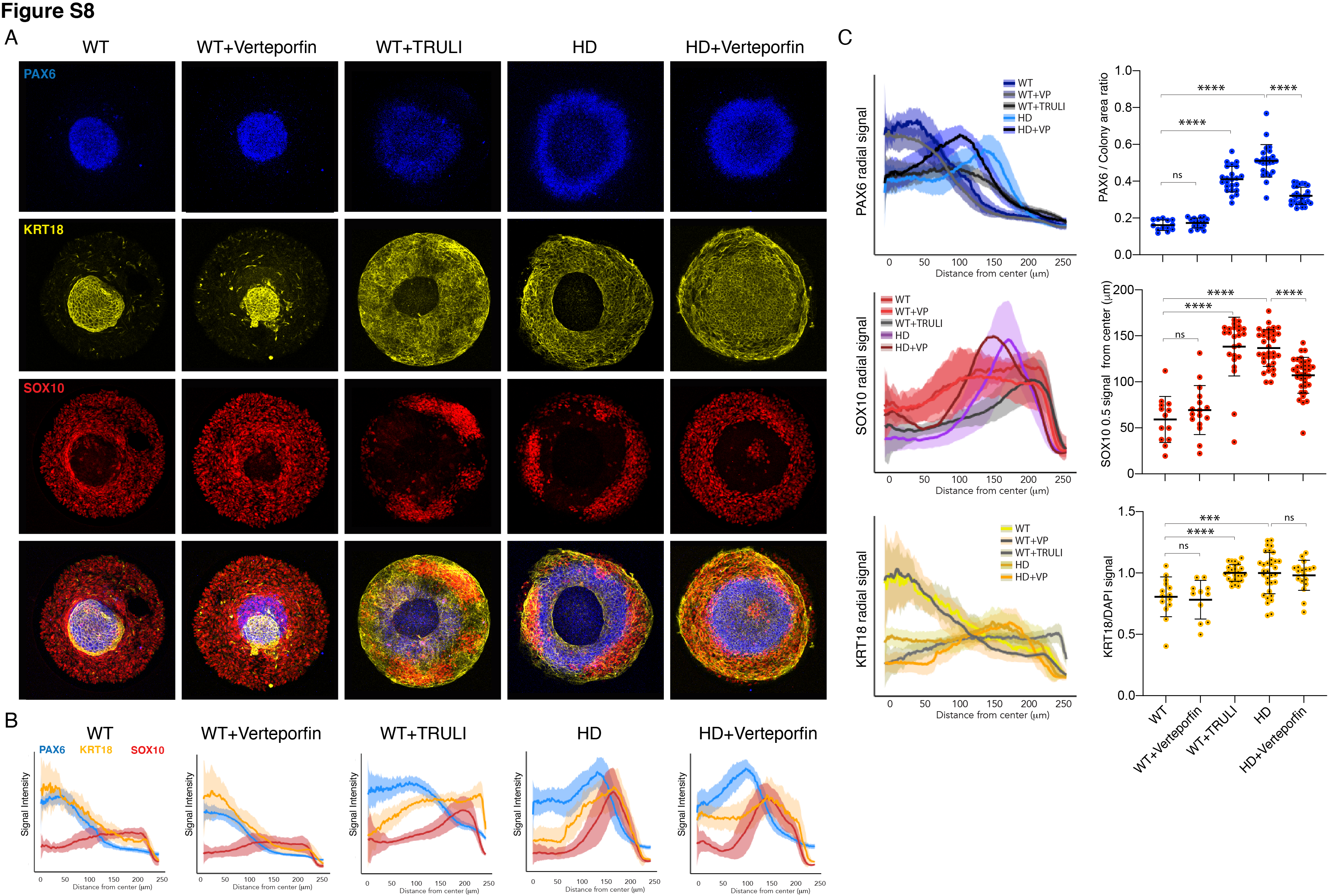

**Figure S8. Perturbation of YAP activity in WT and HD neuruloids.** A) Immunofluorescence images of D7 neuruloids show that treatment with 0.3 µM verteporfin has no effect on WT colonies, whereas treatment with 10 µM TRULI results in the expansion of the central area of NE (PAX6, blue) and enhancement of the epidermal population (KRT18, yellow) at the expense of NC (PAX10, red). This phenotype resembles that of HD colonies. The NE and NC phenotypes of HD colonies are partially rescued by treatment with 0.3 µM verteporfin. B Quantitative analysis of the three experimental conditions shows radial distributions of the lineage markers PAX6, SOX10, and KRT18 in several colonies. C) On the left, radial distributions quantify the lineage markers PAX6, SOX10, and KRT18 in the five experimental conditions. At the right, plots show the area of the PAX6+ central region as a fraction of each entire colony, the radial distance from the colony's center to the half-maximal intensity of the SOX10 domain, and the KRT18 signal normalized by DAPI labeling. ***, *p* < 0.001; ****, *p* < 0.0001; ns, *p* > 0.05 in unpaired *t*‑tests comparing the different experimental conditions.

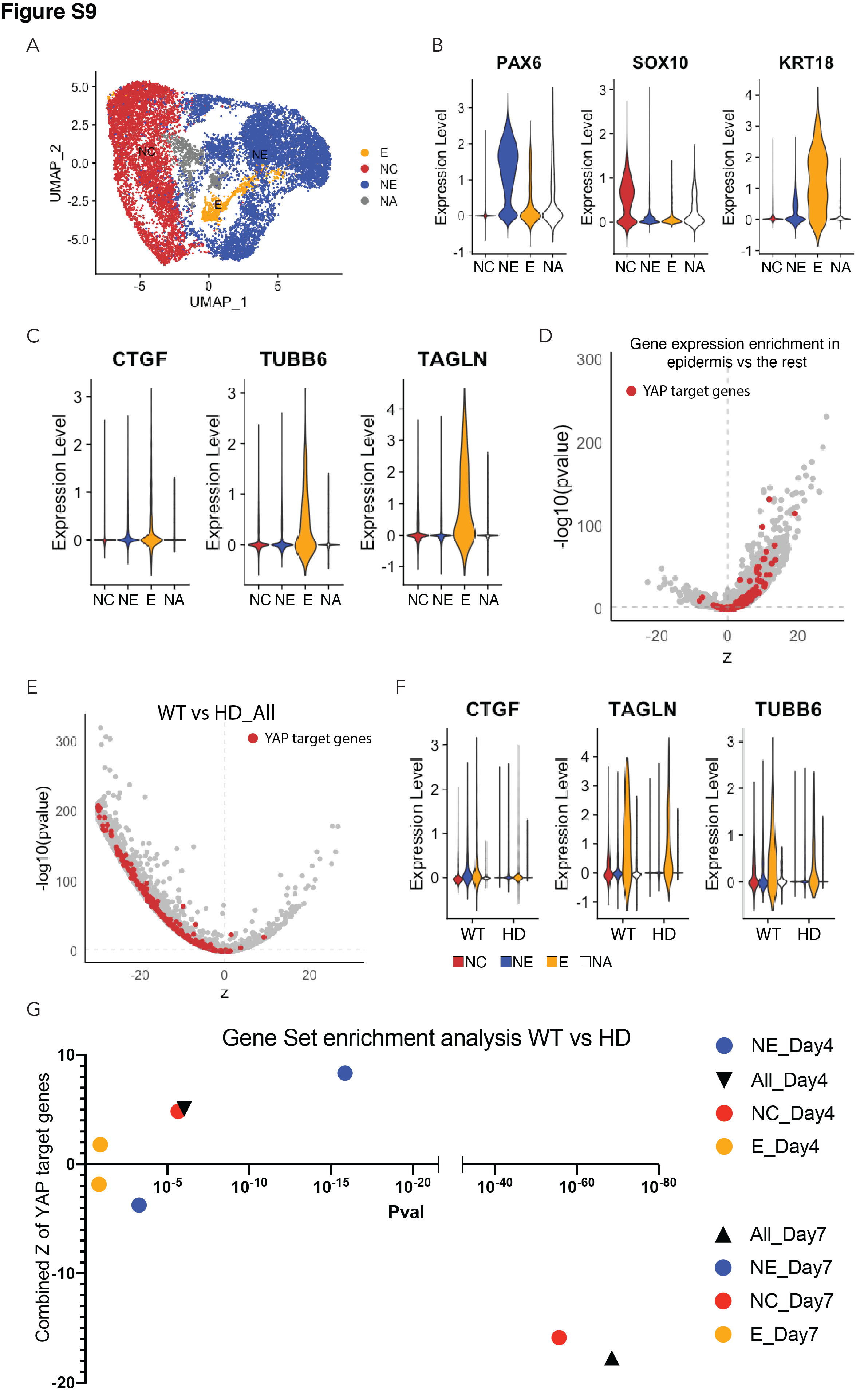

**Figure S9. Expression of YAP target genes in D7 WT and HD neuruloids.** A) Uniform manifold approximation and projection (UMAP) shows the distribution of the three major ectodermal lineages in D7 neuruloid: neural ectoderm (NE, blue), neural crest (NC, red), and epidermis (E, yellow). NA (grey) labels the cell population that was not identified. B) Violin plots quantify the expression of key ectodermal lineage makers: *PAX6* identifies (NE), *SOX10* identifies neural crest (NC), and *KRT18* identifies epidermis (E). C) Violin plots quantify the expression of three representative YAP target genes in the three main ectodermal lineages from scRNA‑seq analysis of D7 neuruloids. D) An analysis of differential gene expression from scRNA‑seq data confirms the greater expression of several YAP target genes (red) in epidermis compared to the remainder of the D7 neuruloid. E) An analysis of differential gene expression from scRNA‑seq data for whole D7 neuruloids shows downregulation of YAP target genes (red) in HD neuruloids with respect to that in WT neuruloids. F) Violin plots show the expression of three representative YAP target genes in the three lineage clusters from D7 WT and HD neuruloids: neural crest (NC, red), neural ectoderm (NE, blue), epidermis (E, yellow) and unidentified (NA, white). G) Gene-set enrichment analysis of YAP target genes reveals reduced YAP activity in D7 HD neuruloids in NE and NC, but not in epidermis. An opposite pattern pertains in less mature D4 neur
